# Coronin1A regulates tumor microenvironment in colitis-associated colorectal cancer in a SUMO-dependent way

**DOI:** 10.64898/2026.02.16.706097

**Authors:** Rohan Babar, Poorvi Saini, Neha Guliya, Vishnu Ashok Kumar E, Prakhar Varshney, Aamir Suhail, Mukesh Singh, Prabhakar Mujagond, Shivangi Tyagi, Lalita Mehra, Dolly Jain, Prasenjit Das, Krishnan Vengadesan, Avinash Bajaj, Vineet Ahuja, Chittur V. Srikanth

**Author notes:** Corresponding Author: C. V. Srikanth. second author with equal contribution (Poorvi Saini, Neha Guliya). third author with equal contribution (Vishnu Ashok Kumar E, Prakhar Varshney).

## Abstract

Inflammatory bowel disease (IBD), comprising Crohn’s disease and Ulcerative colitis, is a group of multifactorial illnesses with persistent gastrointestinal inflammation and a series of undesirable consequences. IBD patients have a three times higher risk of developing colitis-associated colorectal cancer (CAC). A higher mutational burden due to persistent inflammation acts as a driver of dysplasia and even tumorigenesis. While the pathological sequence is known, the molecular mechanisms underlying the transition from chronic inflammation to CAC remain largely elusive. Post-translational modification- SUMOylation plays an integral role in shaping gut inflammation as well as several forms of cancers, including sporadic colorectal cancer. In this study, the contribution of SUMOylation to CAC pathogenesis was characterized. In the AOM-DSS CAC mice model and human IBD patient specimens, SENP5, but not other deSUMOylases, shows an altered expression. Notably, both SENP5 expression dynamics and alterations in the SUMOylome occur in chronically inflamed and neoplastic colon tissues. SENP5 interactome analysis identified Coronin1A (Coro1A), an actin-binding protein predominantly expressed in immune cells. Coro1A also shows a state-specific, distinct expression pattern in the colon. Interestingly, in contrast to wild-type mice, Coro 1A knockout mice were resistant to polyp formation, with reduced cell proliferation, oncogene expression, and epithelial-to-mesenchymal transition (EMT) gene activation, and reduced extracellular matrix (ECM) development and fibrosis, suggesting its integral role in tumorigenesis. WT mice with chronically inflamed colons and polyps have a higher number of M2-like macrophages, with increased abundance of Coro1A, suggesting a role of Coro1A in modulating the tissue microenvironment toward a pro-tumorigenic state. Mechanistically, Coro1A physically interacts with TGF-β RI and regulates TGF-β-TGF-β RI signalling endosome stability, thereby controlling TGF-β-mediated macrophage polarization. Detailed *in vitro* experiments revealed stabilization of Coro 1A through its interaction with SUMOylated Raftlin protein. Overall, Coro 1A is necessary and sufficient for TGF-β signalling, macrophage polarization, and tumorigenesis in CAC.

## Introduction

Colorectal cancer (CRC) is a notorious form of gastrointestinal cancer, with high incidence and alarmingly poor prognosis (Bray et al., 2024; Sung et al., 2025; Schmitt & Greten, 2021). Despite some improvements in therapy, the mortality of CRC remains significantly high, with a 5-year survival rate for stage 4 CRC being less than 10%. Predominant forms of colorectal cancer include (a) sporadic colorectal cancer (hereafter SCRC), which arises due to spontaneous mutation(s) in colorectal epithelium, and (b) colitis-associated colorectal cancer (hereafter CAC), resulting from chronic inflammation in the gastrointestinal tract (Foersch & Neurath, 2014; R. W. Zhou et al., 2023). Multiple routes of tumor initiation and progression in CRC include distinct molecular drivers, such as Adenomatous polyposis coli (APC), TP53, Kirsten rat sarcoma (KRAS), B-Raf protooncogene ser/thre kinase (BRAF), PIK3CA, MMR genes (MSH2, MSH6, and MLH1), and SMAD4, etc (Bogaert & Prenen, 2014; Díaz-Gay et al., 2025; Su et al., 2024). Significantly, the underlying molecular pathology differs across CRC subtypes. SCRC typically follows an adenoma-to-carcinoma sequence driven by prior genetic predispositions (Yamagishi et al., 2016; Yin et al., 2023). Whereas CAC begins with chronic inflammation, progresses through grades of dysplasia, and ultimately ends in carcinoma(Zhou et al., 2023). Consistently, patients with Crohn’s disease (CD) and ulcerative colitis (UC) display a 2-3-fold higher risk of developing CAC (Shawki et al., 2018; Yashiro, 2014; Yin et al., 2023). Chronic inflammation exposes IBD patients to repeated cycles of damage and healing of the mucosa, triggering genetic and/or epigenetic perturbations. These events, referred to as induced field cancerization, prime the tumorigenesis process. Consistent with this, ulcerative colitis patients display a 25-fold higher mutational burden, rendering them susceptible to CAC (Kakiuchi et al., 2020). Infiltrating immune cells and resident fibroblasts of the chronically inflamed gut can induce Wnt-β-Catenin, STAT3/IL-6, and COX2/PGE2 pathways, which are potentially pro-tumorigenic (Dan et al., 2023; Shawki et al., 2018). The tissue microenvironment in this case generates robust reactive oxygen/nitrogen species along with a secretome comprising IL-6, TNF-α, GM-CSF, IL-23, and IL-17, further contributing to DNA damage, epithelial proliferation, and myeloid cell recruitment and activation. Once tumorigenesis is set, the tissue microenvironment is reprogrammed into a tumor-favoring immunosuppressive state. TGF-β, IL-10 loop, or CCR2/CXCL-12 immune evasion pathways induce polarization of immune cells into M2 macrophages, N2 neutrophils, Th2 cells, and T-reg cells (Y. Deng et al., 2025; M. Zhang et al., 2023). Collectively, these events shield dysplastic cells from immune surveillance and foster their proliferation. However, the mechanisms underlying TME reprogramming remain elusive.

Macrophage involvement, characterized as CD11b^+^, F4/80^+^, and Ly6C high and M2 type macrophages, which are referred to as tumor-associated macrophages (TAMs), significantly contributes to the initiation and progression of CAC(Y. Deng et al., 2025; Shin et al., 2023; Hardbower et al., 2017; Y. H. Sheng et al., 2022; Y. Wang et al., 2023; Yuan et al., 2021). Though crosstalk among M2 macrophages, surrounding stromal cells, and other immune cells shapes the tumor microenvironment during CAC progression, there is a lack of mechanistic understanding of these events.

Expansion of immunosuppressive, pro-tumorigenic microenvironment- creates a permissive niche for dysplastic clones to proliferate. A form of cellular post-translational modification (PTM), called SUMOylation, has been shown to be integral in shaping inflammation and cancer. Mediated by the action of three enzymes (E1, E2, and E3), the covalent attachment of SUMO protein isoform (SUMO 1-5) to the target protein takes place, which potentially modulates its function. The deSUMOylase enzymes (SENP1-3 and SENP5-7) then deconjugate the SUMO attachment, making the process reversible. A series of publications have demonstrated a strong connection of SUMOylation with inflammation (Mustfa et al., 2017; Suhail et al., 2019),(Long et al., 2025; Ma et al., 2024). Dysregulation of the SUMOylation pathway has also been linked to EMT and cancer pathophysiology (Gu et al., 2023a; L. Wang et al., 2021). The key molecules involved in the transition from chronic colitis to cancer, such as NF-κB, β-Catenin, p53, AKT, and STAT3, with respect to their activation, stability, and localization, are known to be regulated by SUMOylation. (Gostissa et al., 1999; Lin et al., 2016; Yang et al., 2020; Z. Zhou et al., 2016) Components of the SUMOylation machinery are overexpressed in SCRC and are associated with poor patient prognosis (Du et al., 2016; Liu et al., 2023; Z. Sheng et al., 2025). However, it remains underexplored in CAC to date. Since chronic inflammation in IBD is the primary driver of CAC development, we hypothesized that SUMOylation contributes directly or indirectly to CAC progression. In this study, we demonstrate a dynamic expression pattern of deSUMOylase SENP5 and its prominent interacting partner Coronin1A, an actin-binding immune cell-specific protein, particularly in chronically inflamed regions compared with polyp regions. Coronin1A maintains the canonical TGF-β pathway and thus supports TGF-β-mediated M2 macrophage polarization and modulation of the tumor microenvironment in CAC.

## Results

### Dynamic expression pattern of SENP5 in chronic colitis and colitis-associated colorectal cancer (CAC)

To test the possible role of the SUMOylation machinery in CAC, we employed the previously reported AOM-DSS mice model of CAC (Arnesen et al., 2021). Briefly, 4 mice were included in each group: Control, Chronic colitis (hereafter referred to as CC), and the CAC. After intraperitoneal administration of azoxymethane (AOM) at 10mg/kg to the CAC group, 1% Dextran sulphate sodium (DSS) was given in their drinking water for alternate weeks for 14 weeks to the CAC and CC groups. In contrast, control mice received normal drinking water throughout the experiment. (Fig. 1A). The animals were monitored throughout the experiment, including constant recording of their body weight (Fig. 1B). Post-experimental treatment regimen, the animals were euthanized, and relevant organs were harvested. The successful establishment of the CAC model was evident by the presence of multiple polyps in the distal colon of the CAC group.

**Figure 1.**
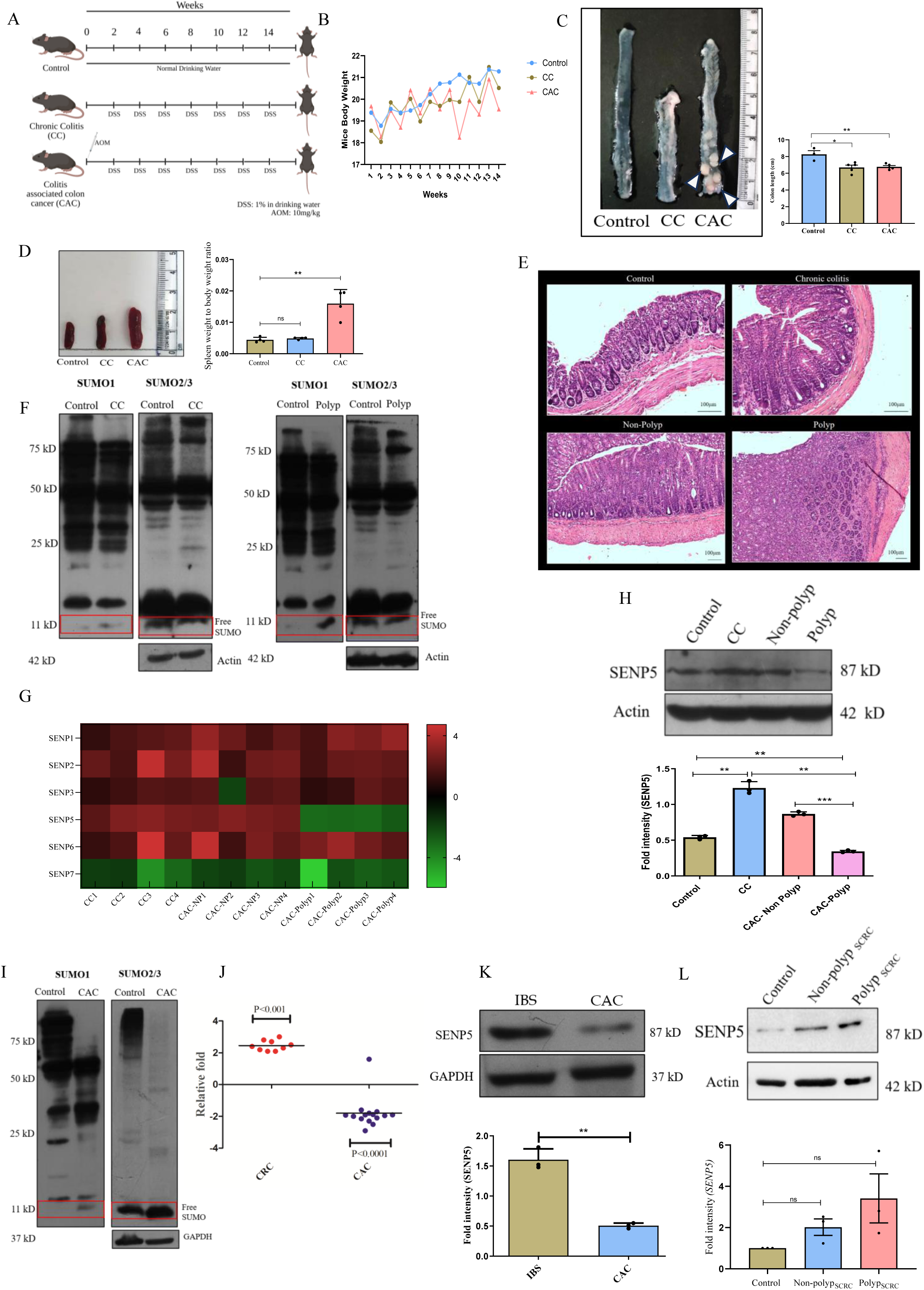
Dynamic expression pattern of SENP5 in chronic colitis and colitis-associated colorectal cancer (CAC) A. Schematic representation of the chemically induced murine CAC model (AOM-DSS) in C57BL/6 mice. B. The graph represents the body weight of experimental animals during the development of the AOM-DSS model. C. A representative image showing the gross morphology of the colon. The graph on the right shows the quantification of colon length. White arrows indicate polyps present at the distal end of the colon. D. A representative image shows the gross morphology of the spleen. The graph to the right shows the spleen weight-to-body weight ratio in the mice. E. Representative Hematoxylin and Eosin-stained colon section from Control, CC, non-polyp, and polyp. F. Immunoblot of SUMO-1 and SUMO-2/3 conjugated proteins (global SUMOylome) from colonic tissue lysates of chronic colitis (CC) and polyps from the murine CAC model, with free SUMO is shown in the red box. G. Heatmap qRT-PCR-based fold change expression of deSUMOylases SENP1, SENP2, SENP3, SENP5, SENP6, and SENP7 in CC, non-polyp, and polyp regions of the CAC murine model relative to the average control values. H. Immunoblot showing SENP5 expression in lysates from colonic tissue of Control, CC, non-polyp, and polyp samples. The graph at the bottom shows the densitometric analysis of the fold intensity of SENP5 expression, normalized to the loading control. I. Immunoblot of SUMO-1 and SUMO-2/3 conjugated proteins (global SUMOylome) from tissue lysates of CAC patient samples with free SUMO is shown in the red box. J. qRT-PCR-based fold change expression of deSUMOylase SENP5 from SCRC (N=9) and CAC (N=15) patient samples relative to the average IBS values (N=10). K. Immunoblot showing SENP5 expression in colonic biopsies from control and CAC patients. The graph at the bottom shows the densitometric analysis of the fold intensity of SENP5 expression, normalized to the loading control. L. Immunoblot showing SENP5 expression in colonic tissue lysates of Control, non-polyp SCRC, and polyp SCRC of Apc^+/+^ and APC^+/-^ model. The graph at the bottom shows the densitometric analysis of the fold intensity of SENP5 expression, normalized to the loading control. Each dot represents (C, D, H, and L) for one mouse, (J) for one patient, and (K) lysate from one patient biopsy. Each box represents one mouse (G). Actin was used as a loading control for F, H, and L, and GAPDH was used as a loading control for I and K. GAPDH was used for gene normalization (G and J). The immunoblots shown here are representative of three biological replicates. Error bars represent mean ±SEM. Statistical analysis performed using the unpaired Student’s t-test, with a p-value <0.05 considered statistically significant. *<0.05, **<0.01, ***<0.001, ****<0.0001, ns: non-significant.

In contrast, the rest of the colon in the CAC (hereafter referred to as Non-polyp) and CC exhibited a pronounced morphological signature of chronic inflammation, characterized by reduced body weight and colon length (Fig. 1 B and C), along with enhanced splenomegaly (Fig. 1D). Histopathology of colon revealed marked infiltration of immune cells along with other signs of chronic inflammation and malignancy in CC and CAC polyps, respectively (Fig. 1E). qRT-PCR-based analysis confirmed significant upregulation of C-Myc in the polyp region (38±5.074), followed by the non-polyp region (22.83±4.642), confirming the development of malignancy in the CAC group. (Fig.S1 A). To test the involvement of SUMOylation in CAC, the abundance of SUMO1 and SUMO2/3 conjugated proteins (hereafter referred to as SUMOylome) in CC and polyp tissue lysates was examined. Compared to the control group, a detectable change in SUMO1 SUMOylome and a pronounced alteration of SUMO2/3 SUMOylome were observed in lysates from both CC and polyps (Fig.1F). An accumulation of free SUMO1 was observed in CC and polyp tissue lysate.

In contrast, no comparable accumulation of SUMO2/3 was observed. To investigate possible involvement of deSUMOylases in this process, qRT-PCR-based expression analysis of deSUMOylases was carried out. Notably, expression of SENP1, SENP2, SENP3, SENP6, and SENP7 remained unchanged, while SENP5 showed a significant increase in expression in the CC (2.61±0.15) and non-polyp regions (2.38±0.25) of CAC. Intriguingly, in stark contrast, SENP5 expression was downregulated in the CAC polyps (-2.8± 0.15) (Fig. 1G). We validated SENP5 expression dynamics in AOM-DSS mice tissue lysate by immunoblotting (Fig. 1H).

Furthermore, the relevance of this finding was investigated in actual CAC patient specimens obtained from the Gastroenterology and Nutrition clinic (AIIMS, Delhi). Biopsies of CAC (N=15), SCRC (N=9), and irritable bowel syndrome (IBS)(N=10) were included in this study. SUMO1 and SUMO2/3 SUMOylome was examined in CAC patient tissue lysates. Consistent with the above findings, a notable alteration in both SUMO1 and SUMO2/3 SUMOylome was observed. The SUMO2/3 SUMOylome was more drastically affected. Moreover, consistent with the AOM-DSS model, increased free sumo was observed in patient tissue lysates (Fig.1I). Furthermore, expression of SENP5 in CAC patient specimens was also downregulated (∼2.06 ± 0.50-fold) (Fig. 1J). Interestingly, in stark contrast, SENP5 was upregulated in sporadic colorectal cancer patient samples (hereafter referred to as SCRC) (Fig. 1J). Immunoblots of SENP5 in CAC patient tissue lysates also revealed reduced expression compared to healthy controls (Fig. 1K).

Additionally, Apc^+/-^ and Apc^+/+^mice were used to develop SCRC (Ren et al., 2019). The animals were administered 2% DSS for 1 week, followed by 14 weeks of incubation to induce tumorigenesis (Fig. S1B). At the conclusion of the experiment, mice were euthanized, and colons were harvested. Prominent polyps were present at the distal end of the Apc^+/-^ mice colon (Fig.S1C). Tissue lysates from the control colon, as well as the polyp and non-polyp regions of Apc^+/-^, were probed for SENP5 by immunoblotting.

Notably, upregulated expression of SENP5 was observed in the polyp region compared to the healthy control and non-polyp region (Fig.1L). Our findings of SENP5 in SCRC patients and the murine model (Apc^+/-^) are consistent with the previous report showing that upregulated SENP5 in SCRC plays a key role in DNA damage repair (Liu et al., 2023). These results clearly indicated a key difference between SCRC and CAC in SENP5 function. The marked difference in SENP5 expression between polyps and non-polyps in the CAC model prompted us to investigate the mechanism in greater detail. To understand the regulatory mechanism driven by SENP5, we conducted SENP5 interactome analysis.

### Altered Interactome of SENP5 in chronic colitis and colitis-associated colorectal cancer (CAC)

The interactome of SENP5 was investigated by performing Co-IP of tissue lysates from control, chronic colitis, non-polyp, and polyp regions of the AOM-DSS mouse model, followed by MS/MS analysis (n=3). An IgG isotype control antibody was used as a negative control. (Fig. 2. A) Pull-down proteins were analyzed by high-resolution tandem mass spectrometry (MS/MS) using a quadrupole time-of-flight (qTOF) mass spectrometer. Initially, 2315 proteins were identified across all four groups; after contaminant removal, the remaining 1,884 proteins were subjected to further analysis. The MaxQuant platform was used to quantify the identified proteins. Peptides found in the isotype control IgG were manually subtracted from the pull-down samples during subsequent analysis. Differential proteome of control, chronic colitis (CC), non-polyp, and polyp obtained from intensity-based absolute quantification illustrated in a Venn diagram (Fig. 2 B), revealing 460 proteins in control, 508 in CC, 510 in non-polyp, and 406 in polyp. Out of which 29 proteins were unique to control, 43 were unique in CC, 74, and 21 were unique in non-polyp and polyp, respectively. Notably, 282 proteins are present in all four conditions. In the identified set of proteins, we detected SUMO2. Recent structural studies have demonstrated the active, specific interaction between SENP5 and SUMO2, underscoring the robustness of our interactome approach. (Sánchez-Alba et al., 2025) A list of proteins identified across all four conditions is provided in the supplementary.

**Figure 2.**
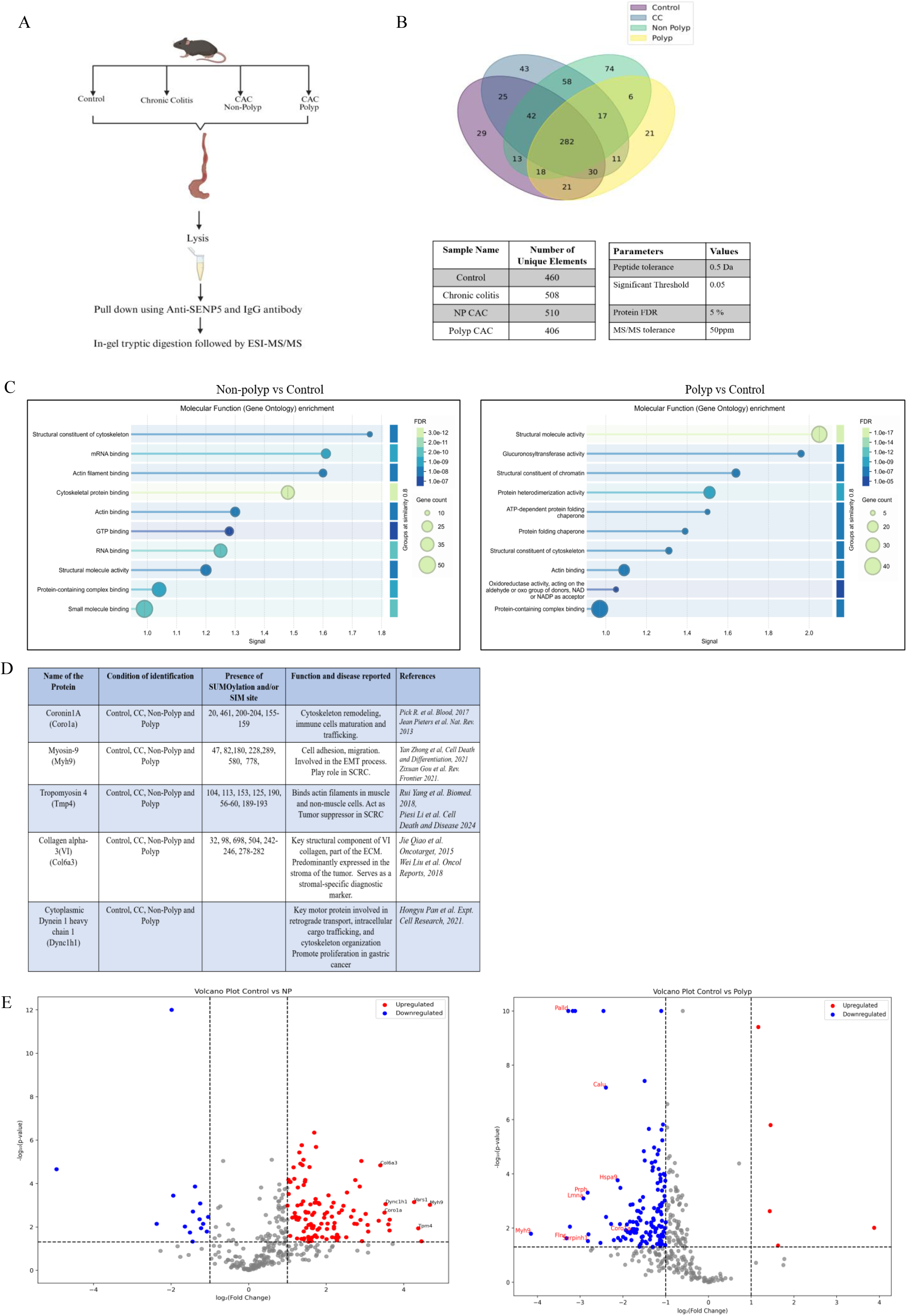
Altered interactome of SENP5 in chronic colitis and colitis-associated colorectal cancer. A. Schematic representation of SENP5 pull-down using anti-SENP5 antibody from Control, CC, non-polyp, and polyp tissue lysates. Isotype antibody (IgG) was used as a negative control. B. Venn diagram showing the number of unique and common proteins across the Control, CC, non-polyp, and polyp groups. C. The Gene Ontology (GO) Pathway analysis (molecular function) of differentially enriched SENP5 interacting proteins in the non-polyp and polyp region, as compared to the Control. Pathway enrichment significance was assessed using hypergeometric and multiple test correction at FDR less than 0.05. D. Table of shortlisted potential SENP5 interacting proteins from the Actin binding function pathway based on their functional role in cancer or any disease progression, and having a SUMO binding or interacting motif. E. Volcano plot of differentially enriched proteins with SENP5 in non-polyp and polyp as compared to the Control, with shortlisted potential SENP5 interactors marked. Here, significance is defined as fold change >2 and p-value < 0.05.

Differential enrichment analysis of SENP5-interacting proteins showed a distinct pattern across all four conditions. 114 proteins were enriched, and 15 proteins were depleted in their interaction with SENP5 significantly in the non-polyp region. Under CC,39 proteins were enriched with SENP5, and 13 proteins were depleted in their interactions with SENP5.

In the polyp group, 132 proteins showed reduced interaction with SENP5, and five proteins showed enrichment with SENP5. Here, the definition of significance is a fold change greater than two and a *p*-value <0.05.

To understand the biological significance of proteins differentially enriched for SENP5, Gene Ontology (GO) pathway analysis was performed on enriched and depleted proteins across all three disease conditions. Pathway enrichment significance was assessed using hypergeometric and multiple test correction at FDR less than 0.05. The *p*-values are corrected for multiple testing within each category using the Benjamini-Hochberg procedure. As both non-polyp and chronic colitis represent states of chronic inflammation, we chose non-polyp samples for differential GO pathway enrichment analysis. In this enrichment analysis, polyp and non-polyp samples were independently compared with the control to identify pathways associated with the tumor and the adjacent non-tumor area.

Molecular function and cellular component-based GO pathway analysis revealed a profound enrichment of SENP5-interacting proteins with actin filament binding or actin binding functions, and the actin cytoskeleton cellular component, particularly in the non-polyp region. While the enrichment of these proteins was very feeble in the polyp region (Fig. 2C) (Fig.S2. A). These results suggest that SENP5 displays a more pronounced interaction with proteins involved in Actin binding or linked to the actin cytoskeleton in the non-polyp region. In contrast, such interactions are markedly reduced in polyp regions. From the significantly enriched list of proteins falling under the Actin binding or Actin filament binding function, we shortlisted proteins Coro1A, Tmp4, Myh9, Col6A3, and Dync1h1, based on their roles in cancer or disease progression, as well as the presence of prominent SUMO-binding or SUMO-interacting sites (Fig.2D). The differential enrichment of selected genes was visualized on volcano plot, revealing their prominent enrichment in the non-polyp and CC region and abolished enrichment in polyp region. (Fig 2. E) (Fig.S2.B) It underlines a shift in the interactome of SENP5 from the non-polyp to the polyp region, indicating a crucial role of these proteins in disease progression. Among the selected candidates, Coro1A (Coronin 1A) stood out as the most prominent pick. The immune cell-specific protein Coro1A is known to interact with filamentous Actin and regulate actin dynamics by regulating the Arp2/3 complex (Galkin et al., 2008; Humphries et al., 2002) (Machesky et al., 1997; Shatery Nejad et al., 2025). Coronin 1A plays a role in T-cell maturation, as well as T-cell receptor-based signaling, neutrophil trafficking, and immune deficiency (Castro-Castro et al., 2011; Mueller et al., 2008; Pick et al., 2017).Additionally, there is multiple evidence of Coronin1A involvement in immunological pathways associated with chronic inflammation and its related manifestations, such as lung fibrosis, lupus nephritis, and non-alcoholic steatohepatitis (NASH), and autoimmune diseases such as systemic lupus erythematosus and multiple sclerosis. (X. Deng et al., 2024; Nicolaou et al., 2020; Siegmund et al., 2016) (Haraldsson et al., n.d.)In light of these, we set forth to investigate the role of Coronin-1A in CAC progression.

### Coro1A is an interactor of SENP5

The interactome analysis enabled us to identify Coronin-1A (hereafter referred to as Coro1A) as an interactor of SENP5. Coro1A-SENP5 interaction was scrutinized using a co-immunoprecipitation (Co-IP) assay. The immunoblot clearly showed strong binding of Coro1A to SENP5, particularly in tissues from CC and non-polyp areas of CAC mice (Fig. 3A). Furthermore, the expression profile of Coro1A was examined in the AOM-DSS mouse model. An elevated expression of Coro1A in CC and, interestingly, in the non-polyp region of CAC was observed. In contrast, in the polyp, much lower expression was detected (Fig. 3B). These results were validated in a murine colitis genetic model using IL10^-/-^ mice. IL10^⁻/⁻^ mice spontaneously develop colitis (Gunasekera et al., 2020).To establish a synchronized CAC in this background, the animals were administered AOM (10mg/kg) intraperitoneally, followed by 7 alternating cycles of low-dose 1% DSS (Fig. S3A). Post-treatment, the mice were euthanized, and the relevant organs were harvested. Notably, we did not detect any polyps in CAC mice, despite probing the colons of CC and CAC mice for Coro1A expression. Immunoblotting of colon lysates revealed a pronounced increase in Coro1A expression in both CC and CAC tissues compared to untreated controls (Fig. 3C), confirming that Coro1A upregulation is not restricted to a single chronic colitis model but is found in a broad range of chronic colitis models. Next, the expression of Coro1A was analyzed in an acute colitis mouse model. The acute colitis model was developed as described by (Suhail et al., 2019)(Fig. S3B). Coro1A expression was not modulated in the acute colitis mouse model (Fig. S3C). These results implied that Coro1A induction is limited to chronic colitis. Additionally, we probed Coro1A in colonic lysates from the Apc^+/-^ SCRC model and found no significant differences in its expression; however, a trend of decreasing Coro1A expression was observed in the non-tumor adjacent area and the tumor area. (Fig. S3D). Next, the relevance of Coro1A expression in chronic gut inflammation was examined in human colon biopsies. Biopsy samples from UC (N=5) patients also showed significantly elevated Coro1A expression compared to those from IBS patients (N=4), as shown by immunoblot (Fig. 3D). These findings suggest a potential role for Coro1A in the development of colitis-associated cancer.

**Figure 3.**
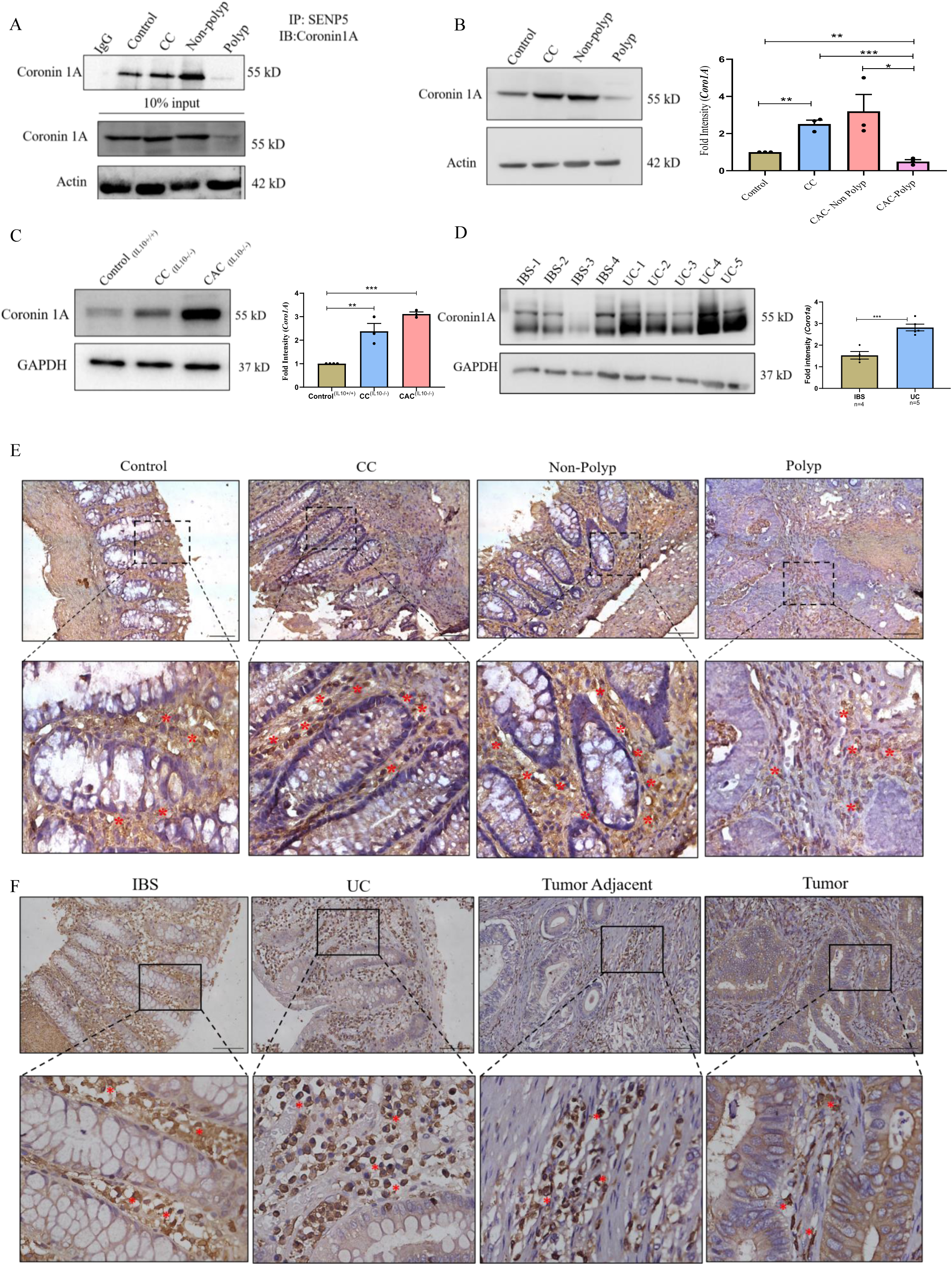
Coro1A as an interactor of SENP5. A. Top: Colonic tissue lysates of Control, CC, non-polyp, and polyp were immunoprecipitated with anti-SENP5 antibody and immunoblotted for Coro1A. Bottom: 10% of input loaded from the same tissue lysate and immunoblotted for Coro1A and Actin. Isotype antibody (IgG) was used as a negative control. B. Immunoblot showing Coro1A expression in colonic tissue lysates of Control, CC, non-polyp, and polyp. The graph on the right shows densitometric analysis of fold intensity of Coro1A expression, normalized to loading control. C. Immunoblot showing Coro1A expression in colonic tissue lysates from Control, CC, and CAC colon of an AOM-DSS model performed in IL-10^-/-^ mice. The graph on the right shows densitometric analysis of fold intensity of Coro1A expression, normalized to loading control. D. Immunoblot showing Coro1A expression in human colonic biopsies of IBS (N=4) and ulcerative colitis (N=5) patients. The graph on the right shows densitometric analysis of fold intensity of Coro1A expression, normalized to loading control. E. Immunohistochemistry of Coro1A in colonic tissue sections of Control, CC, non-polyp, and polyp region of AOM-DSS mice model. Scale bar 100μm. F. Immunohistochemistry of Coro1A in archived colonic biopsy slides from IBS, UC, and CAC patients. Scale bar 100μm. Each dot represents (B and C) one mouse or (D) one patient biopsy tissue lysate. Actin was used as a loading control (A and B), and GAPDH was used as a loading Control (C and D). The inset shows a zoomed image of a colon section. Asterisks show Coro1a expression (E and F). The immunoblots shown here are representative of three biological replicates. Error bars represent mean ±SEM. Statistical analysis performed using the unpaired Student’s t-test, with a p-value <0.05 considered statistically significant. *<0.05, **<0.01, ***< 0.001, ****<0.0001, ns: non-significant.

Together, these findings indicate that Coro1A upregulation is a feature specific to chronic inflammatory settings and inflammation-driven tumorigenesis, rather than sporadic tumor development. To gain insights into the subcellular localization of Coro1A, immunohistochemistry (IHC) was performed. Enhanced expression of Coro1A was observed in infiltrating immune cells in the CC and non-polyp areas compared with the control. Coro1A protein was detected in the polyp region, though with less intensity as compared to CC and non-polyp (Fig. 3E). Furthermore, these observations were confirmed in patient biopsy-derived archived slides, which showed a consistent increase in Coro1A expression in infiltrating immune cells in UC and the adjacent inflamed CAC tumor area compared to the IBS control. Notably, a prominent presence of Coro1A in the CAC tumor, albeit less than in non-polyp and UC, was observed (Fig. 3F). These results reveal that Coro1A is uniformly present in immune cells and is consistently upregulated during chronic inflammation. However, an unusually high abundance of Coro1A in the non-polyp region of CAC prompted us to investigate its role in CAC development.

### Absence of Coro1A confers resistance to CAC development in mice

To understand the relevance of increased Coro1A expression in infiltrating immune cells, we employed Coro1A KO mice (hereafter, Coro1A^KO^). Coro1A ^KO^ mice were generated in the laboratory of J.M. Penninger (IMBA, Vienna) by using *a Cre/LoxP-*based gene-targeted strategy. The deleted region encompasses the entire coding sequence after exon 1 (Kaminski et al., 2011). Complete KO of the Coro1A gene was confirmed by immunoblot of Coro1A from colonic tissue lysates of Wild-type mice (WT) and Coro1A^KO^ mice (Fig. S4A and B).

To understand the relevance of Coro1A in CAC development, WT and Coro1A^KO^ were subjected to the AOM-DSS treatment as described in Fig.1A. A schematic representation of the experimental design, which lasted for 14 weeks, is shown in Fig. 4A. The animals were monitored throughout the experiment, including constant recording of their body weight (Fig. S4C). Post experiment, mice were euthanized, and the colon, spleen, and other relevant organs were harvested. Gross morphological parameters were evaluated to compare the extent of disease development in WT and Coro1A^KO^ mice. The length of the colon and splenomegaly were analyzed in both CC and CAC groups across both genotypes. A considerable reduction in colon length of CC and CAC categories of each genotype was observed, indicating the establishment of colitis (Fig. S4.D). No significant differences were observed in the respective sub-categories between the WT and Coro1A^KO^ groups. However, the colon from WT mice in the CAC group showed a wider distal end than those in all other groups from both genotypes, suggesting the possible presence of polyps. In line with this, the colon from WT mice in the CAC group showed the development of multiple polyps, a feature absent in Coro1A^KO^ mice (Fig. 4B). Moreover, there was an evident splenomegaly in the case of the CAC^WT^ group, which was more pronounced in the WT as compared to their KO counterparts (Fig. S4E). H&E staining of colonic tissue sections from WT and Coro1A^KO^ mice across all conditions was carried out. Sections from CC^WT^ and CAC ^WT^ groups showed prominent markers of colitis and cancer, including high-grade dysplasia, mild field effect, collagen deposition, distorted crypt architecture, a widened muscularis layer, and moderate inflammation with immune cell infiltration.

**Figure 4.**
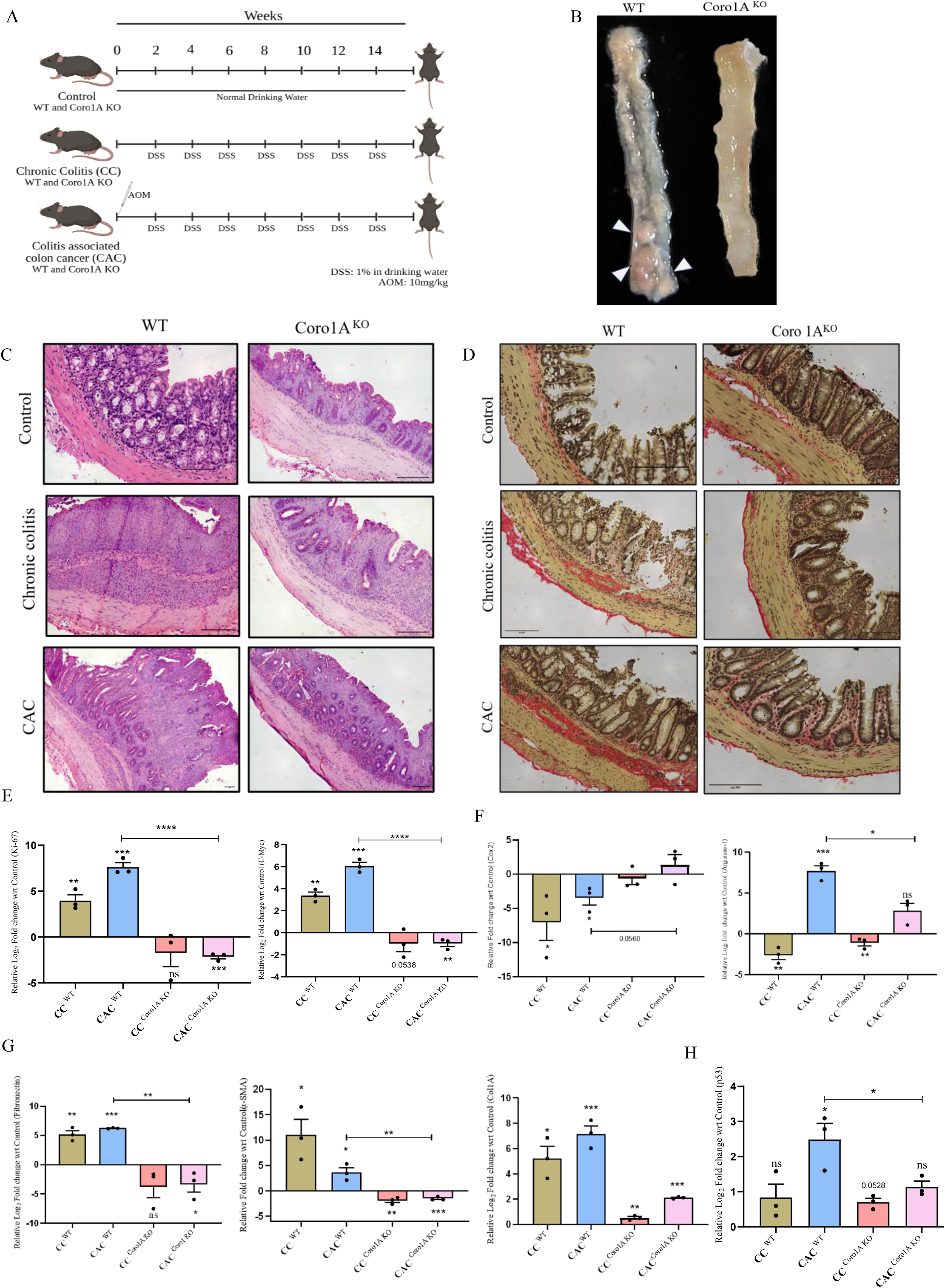
Absence of Coro1A confers resistance to CAC development in mice: A. Schematic representation of the chemically induced murine CAC model (AOM-DSS) in WT (C57BL/6) and Coro1 A ^KO^ (C57BL/6) mice. B. Representative longitudinal cut-open colon image from the CAC group of WT and Coro1A ^KO^ mice. White arrows indicate polyps present at the distal end of the WT colon. C. Representative Hematoxylin and Eosin-stained colon section from Control, CC, and CAC group of WT and Coro1A ^KO^ mice. D. Representative Sirius Red-stained colon section from Control, CC, and CAC group of WT and Coro1A^KO^ mice. E. qRT-PCR-based fold change expression of Proliferation and Oncogene; Ki67 and C-Myc relative to the average control values. F. qRT-PCR-based fold change expression of pro-inflammatory and anti-inflammatory markers, COX-2 and Arginase-1, relative to the average control values. G. qRT-PCR-based fold change expression of ECM markers, Fibronectin, α-SMA, and Col1A1 relative to the average control values. H. qRT-PCR-based fold change expression of p53 relative to the average control values. Each dot represents one of the mice (E, F, G, and H). Samples analyzed by qRT-PCR include CC ^WT^, CAC ^WT^, CC ^Coro1A^ ^KO^, and CAC ^Coro1A^ ^KO^. GAPDH was used for Gene normalization (E, F, G, and H). Error bars represent mean ± SEM. Statistical analysis performed using the unpaired Student’s t-test, with a p-value <0.05 considered statistically significant. *<0.05, **<0.01, ***< 0.001, ****<0.0001, ns: non-significant.

In contrast, the CC and CAC groups of Coro1A^KO^ mice exhibited no or very low-grade dysplasia, field effect, crypt distortion, and collagen deposition, accompanied by prominent immune cell infiltration and inflammation, as indicated by the histopathology score (Fig. 4C and Fig. S4F). Further Sirius red staining showed prominent collagen deposition in CC^WT^ and CAC ^WT.^In contrast, Coro1A^KO^ mice exhibited strikingly less collagen deposition in CC and CAC, indicating attenuated extracellular matrix (ECM) deposition in Coro1A ^KO^ mice compared with WT. (Fig.4D). The lack of polyp formation and absence of adenocarcinoma in Coro1A^KO^ CAC mice necessitated the investigation of molecular markers associated with proliferation, inflammation, extracellular matrix (ECM) deposition, and epithelial-to-mesenchymal transition (EMT) in colonic tissue lysates from both genotypes across all samples.

Both Ki-67, a proliferation marker, and C-Myc, a protooncogene, were markedly elevated in CC^WT^(2.973±0.6296; 2.363±0.3234) and CAC^WT^ (6.600±0.05003; 5.050±0.3396) mice, whereas they were downregulated in the CC^Coro1aKO^(-2.723±1.513;-1.987±0.7342) and CAC^Coro1aKO^(-3.137±0.2270; -1.980±0.2722) counterparts (Fig.4.E). Expression of E-Cadherin was dramatically downregulated in CC^WT^(-29.10 ± 9.570) while the CAC^WT^ group showed marked upregulation (14.22±1.650). Interestingly, both features were completely absent in the Coro1AKO groups. This pattern of E-cadherin expression aligns with recent reports of elevated E-cadherin expression in solid tumors (Burandt et al., 2021; Rodriguez et al., 2012) (Fig. S4G). Cox2, a pro-inflammatory marker, was downregulated significantly in CC(-8.043±2.687) and CAC(-4.457±1.057) of WT, whereas Coro1A^KO^ CC and CAC showed very subtle changes. Anti-inflammatory and pro-wound healing marker-Arginase-1(Arg1) was markedly upregulated in CAC^WT^(6.700±0.6255), and downregulated in CC^WT^(-3.600±0.5615). In the Coro1A^KO^ CC and CAC mice, this pattern was observed, but with significantly lower values (-2.097±0.3783; 1.847±0.8995) (Fig.4F). Together, these results indicate a tendency toward higher cell proliferation, higher oncogenic, excessive wound healing, anti-inflammatory, and lower pro-inflammatory activity in WT mice; however, these features were either absent or diminished in Coro1A^KO^ mice.

Further analysis of ECM remodeling genes, including Fibronectin, α-SMA, and Col1A1, revealed significant upregulation in WT mice from the CC(4.183±0.6451; 10.04±3.008; 4.227±0.9239) and CAC(5.233±0.04910; 2.633±0.9347; 6.157±0.6388) groups. In contrast, their expression was comparatively lower in respective groups of CC^Coro1A^ ^KO^ (-4.727±1.916; -2.898± 0.3824; 0.5067± 0.1074) and CAC^Coro1AKO^(-4.370±1.344;-2.513±0.1995;1.100±0.04933) mice (Fig.4G). Consistently, EMT-associated transcription factors SNAIL and SLUG were also upregulated in CAC ^WT^ (7.072±1.964; 24.33± 5.503) relative to the counterpart Coro1A^KO^ tissues group (-7.223± 0.2706; 0.5819± 0.3946) (Fig.S4H). Additionally, p53 was upregulated in CAC^WT^ (1.490± 0.4539) compared to CC^WT^ (-0.1667±0.3820) and both Coro1A KO conditions(-0.300±0.1102; 0.1367±0.1637). It is consistent with reports of p53 overexpression in UC, CAC, and sporadic colorectal cancer (Khursheed et al., 2024; Lu et al., 2017; Popp et al., 2016) (Fig. 4H). Altogether, these findings demonstrate that molecular footprints associated with proliferation, inflammation, ECM remodeling, and EMT are pronounced in WT mice, correlating with their increased susceptibility to CAC development compared to Coro1A^KO^ mice.

### Increased expression of Coro1A is associated with M2 macrophages

To understand the immune cell type that shows enhanced Coro1A expression in chronic inflammatory conditions and plays a crucial role in CAC development, we analyzed immune cells isolated from the lamina propria of control, CC, non-polyp, and polyp tissues from WT mice by flow cytometry. Flow cytometry was performed using predefined gating strategy, as shown in Fig. S5A. Analysis revealed, in contrast to the control group, a marked increase in the number of F4/80^+^ and CD11b^+^ macrophages in CC, non-polyp, and polyp regions (Fig. 5A and B), while the neutrophil count (CD45^+^ and Ly6G^+^) was decreased in the CC and non-polyp regions, but unexpectedly elevated in the polyp tissue (Fig S.5 B and C). To confirm whether Coro1A is increased in infiltrating macrophages, the Median fluorescence intensity (MFI) of Coro1A during flow cytometric analysis was calculated. MFI analysis revealed increased abundance of Coro1A in infiltrating macrophages across all three conditions (Fig. 5C). While decreased abundance of Coro1A was observed in neutrophils in the CC and non-polyp regions, it was strikingly elevated in the polyp region (Fig. S5D). Under prolonged chronic inflammation, dynamic macrophages exhibit remarkable plasticity, adapting to either a pro-inflammatory (M1) or an anti-inflammatory (M2) state. Overall, macrophages, particularly anti-inflammatory, pro-tumorigenic M2 macrophages, utilize a variety of signaling cascades to promote CAC. (Hardbower et al., 2017; Wang et al., 2023; Yuan et al., 2021 Suarez-Lopez et al., 2018) Further analysis demonstrated a gradual enrichment of(Arg^+^-1) M2-like macrophages in CC, non-polyp, and polyp regions compared to the iNOS^+^ M1-like macrophages (Fig. 5D). A significantly higher M2 (Arg^+^) /M1 (iNOS^+^) ratio was seen in the area of non-polyp and polyp (Fig.5E). Subsequent analysis revealed increased MFI of Coro1A is associated with Arg^+^ M2 macrophages (Fig. 5F), whereas iNOS^+^ M1 macrophages did not show such a trend (Fig. S5E).

**Figure 5.**
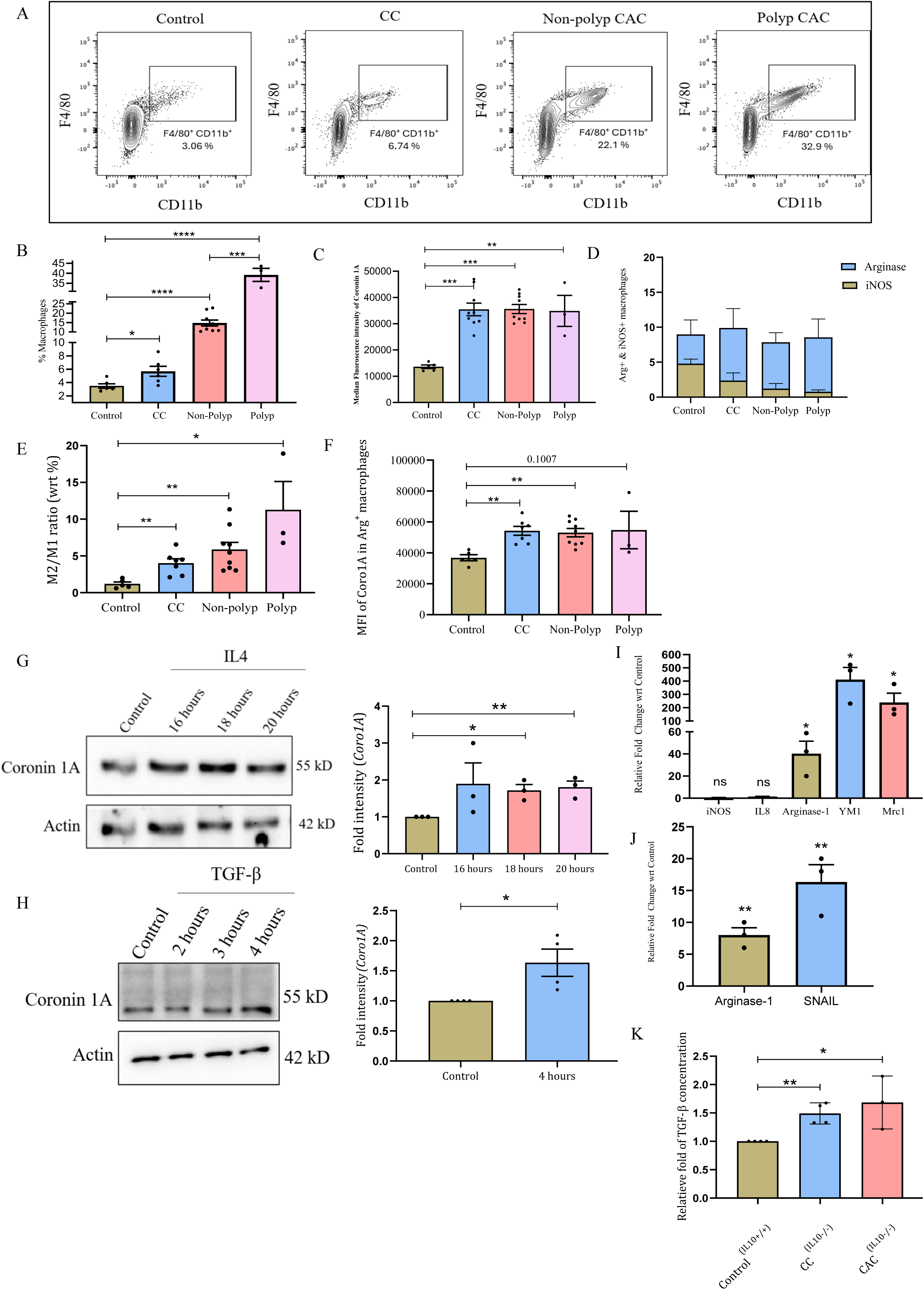
Increased expression of Coro 1A is associated with M2 macrophages. A. Representative counter plots showing F4/80^+^ and CD11b^+^ macrophage infiltration in Control, CC, non-polyp, and polyp regions. B. Percentage infiltration of macrophages in Control, CC, non-polyp, and polyp as determined by flow cytometry. C. Median Fluorescence intensity (MFI) of Coro1A in infiltrated macrophages across Control, CC, non-polyp, and polyp as measured by flow cytometry. D. Representative plot showing the distribution of iNOS^+^ (%) and Arg-1^+^ (%) macrophages across Control, CC, non-polyp, and polyp. E. Ratio of M2 (Arg-1^+^) to M1 (iNOS^+^) macrophages in all four conditions: Control, CC, non-polyp, and polyp, as determined by flow cytometry. F. MFI of Coro1A in Arg-1^+^ macrophages across Control, CC, non-polyp, and polyp. G. Immunoblot showing expression of Coro1A in bone-marrow-derived macrophages (BMDMs) cells treated with IL-4 for 16, 18, and 20 hours. The graph on the right shows the densitometric analysis of the fold intensity of Coro1A expression, normalized to the loading control. H. Immunoblot showing expression of Coro1A in RAW 264.7 cells treated with TGF-β for 2,3, and 4 hours. The graph on the right shows the densitometric analysis of the fold intensity of Coro1A expression, normalized to the loading control. I. qRT-PCR analysis showing fold-change expression of M1(iNOS and IL-8) and M2 (Arg-1, YM1, and Mrc1) macrophage markers in BMDMs treated with IL-4. J. qRT-PCR analysis showing fold-change expression of M2 macrophage markers, Arginase-1, and SNAIL in RAW264.7 cells treated with TGF-β. K. Quantification of TGF-β levels in colonic tissue lysates from AOM-DSS mice, measured by ELISA. Each dot represents (B, C, E, F, and K) one mouse, or (G, H, I, and J) independent experiments of 3 biological replicates. Actin is used as a loading control (G and H). GAPDH used for Gene normalization (I and J). The immunoblots shown here are representative of three biological replicates. Error bars represent mean ± SEM. Statistical analysis performed using the unpaired Student’s t-test, with a p-value <0.05 considered statistically significant. *<0.05, **<0.01, ***< 0.001, ****<0.0001, ns: non significant.

To gain a mechanistic understanding of these observations, *in vitro* experiments were conducted. Macrophages were treated with IL-4 and TGF-β to induce an M2-like phenotype as previously reported (Stein et al., n.d.)(Zhang et al., 2016).Notably, exposure to these two cytokines increased intracellular Coro1A levels (Fig. 5 G and H). This phenotype was accompanied by the upregulation of several canonical M2 macrophage markers 14 hours post cytokine treatment (p.c.t) (Fig. 5I and J).

Considering TGF-β being one of the inducers of Coro1A, the levels of TGF-β in the tissue lysates of the AOM-DSS murine model were verified. Found an increased abundance of TGF-β in the CC and CAC colon of IL-10^-/-^ AOM-DSS mice, as determined by ELISA, underscoring the connection between TGF-β and Coro1A signaling (Fig. 5 K). Collectively, these results suggest that Coro1A is involved in the function of M2-like macrophages and thus modulates the tumor microenvironment.

### Coro1A interacts directly with the TGF-βRI and controls TGF-β-mediated M2 macrophage polarization

Previous reports have implicated Coro1A in regulating TGF-βRI and in the differentiation of Th17 CD4^+^ T cells (Kaminski et al., 2011). Based on this, we set forth to investigate the possible mechanism by which Coro1A crosstalk with TGF-β-signaling in macrophages. Co-IP analyzed the potential interaction between Coro1A and TGF-βRI. RAW264.7 cell lysate was pulled down with an anti-Coro1A antibody, followed by immunoblotting against TGF-βRI. A direct interaction of Coro1A with TGF-βRI was evident (Fig. 6.A). Furthermore, structured illumination microscopy (SIM) revealed a direct interaction between TGF-βRI and Coro1A, as evidenced by strong colocalization signals (Fig.6B). Furthermore, interaction of Coro1A was not restricted to TGF-β RI; it also interacted with TGF-β RII (Fig. S. 6A).

**Figure 6.**
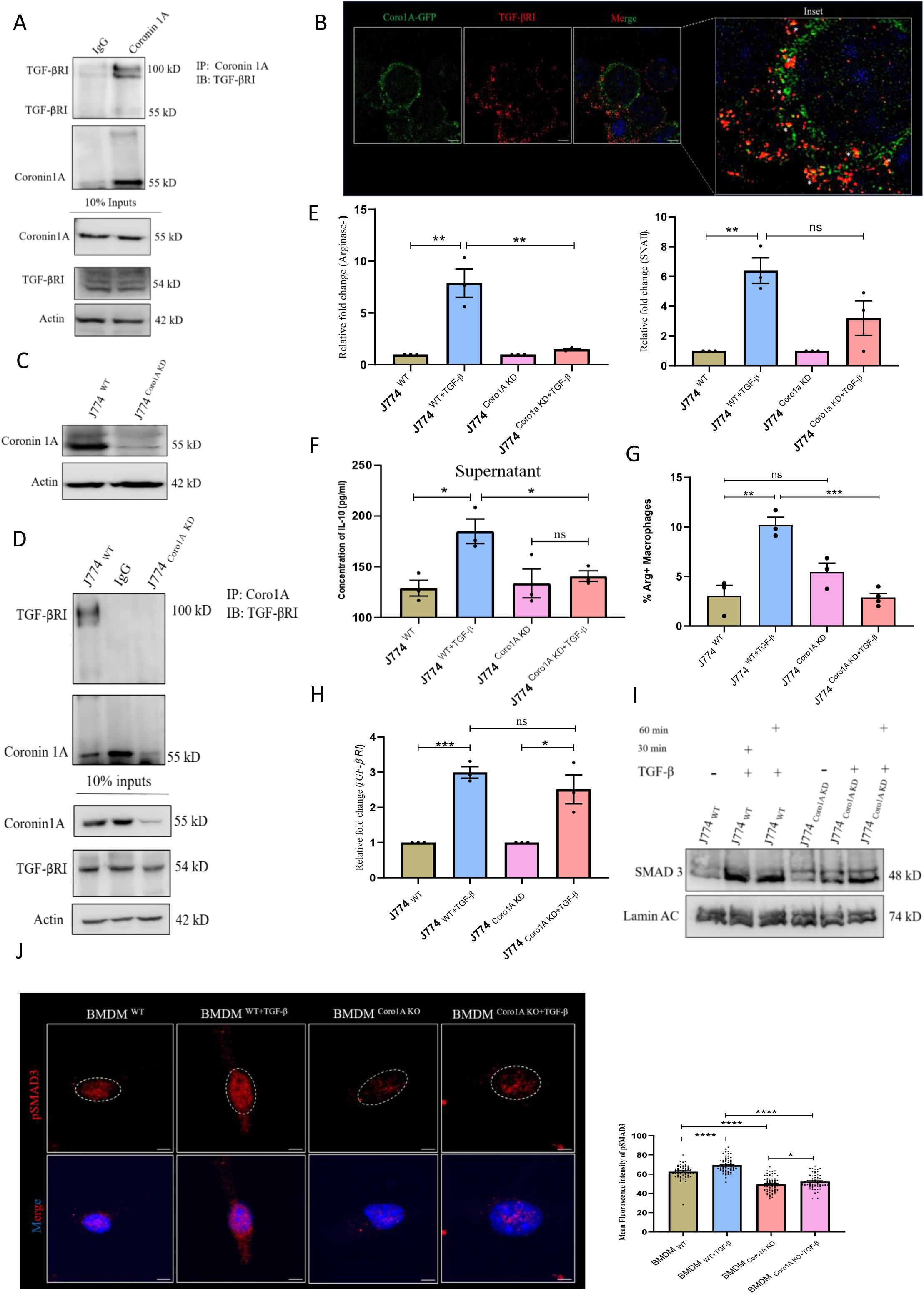
Coro1A interacts with TGF-β receptors directly and controls TGF-β-mediated M2 macrophage polarization. A. Top: RAW264.7 cell lysates immunoprecipitated with anti-Coro1A antibody and immunoblotted for TGF-βRI. Bottom:10% of input loaded from the same cell lysate and immunoblotted for Coro1A, TGF-βRI, and Actin. Isotype antibodies (IgG) were used as a negative control. B. Representative structured illumination microscopy (SIM) showing cellular localization of TGF-βRI (Red) and Coro 1A (Green). The asterisks in the zoomed version of the merged image show colocalization of TGF-βRI (Red) and Coro1A-GFP (Green). C. Immunoblot confirming loss of Coro1A protein expression in stable J774 ^Coro1A^ ^KD^ cells compared to J774^WT^ cells. D. Top: J774 ^WT^ and J774 ^Coro1A^ ^KD^ cell lysates immunoprecipitated with anti-Coro1A antibody and immunoblotted for TGF-βRI. Bottom: 10% of input loaded from cell lysate and immunoblotted for Coro1A, TGF-βRI, and Actin. Isotype antibodies (IgG) were used as a negative control. E. qRT-PCR-based fold-change expression of Arginase-1 (left) and SNAIL (right) in J774 ^WT^ and J774 ^Coro1A^ ^KD^ cells with and without TGF-β stimulation. F. Concentration analysis of IL-10 in the cell culture supernatant of J774 ^WT^ and J774^Coro1A^ ^KD^ cells with and without TGF-β stimulation using ELISA. G. Flow cytometry-based analysis of Arginase-1 positive cells in J774 ^WT^ and J774 ^Coro1A^ ^KD^ cells with and without TGF-β stimulation. H. qRT-PCR-based fold-change expression of TGF-βRI in J774 ^WT^ and J774 ^Coro1A^ ^KD^ cells with and without TGF-β stimulation. I. Immunoblot showing nuclear accumulation of SMAD3 post 30 and 60 minutes of TGF-β exposure to J774 ^WT^ and J774 ^Coro1A^ ^KD^ cells. J. Representative confocal images of BMDM ^WT^ and BMDM ^Coro1A^ ^KO^ cells with and without TGF-β stimulation, stained for pSMAD3 (Red). A dotted white circle demarks nuclear boundaries. The graph to the right illustrates the mean fluorescence intensity of pSMAD3 in the nuclei (n = 60). (Scale bar = 5μm). Each dot represents (E, F, G, and H) one independent experiment of a biological experiment, and (J) one cell. Actin was used as a loading control (A, C, and D). Lamina AC was used as a loading control (I), GAPDH was used for Gene normalization (E and H). The immunoblots shown here are representative of three biological replicates. Error bars represent mean ± SEM. Statistical analysis performed using the unpaired Student’s t-test, with a *p*-value <0.05 considered statistically significant. *<0.05, **<0.01, ***<0.001, ****<0.0001, ns: non-significant.

The role of Coro1A in macrophage function was next assessed. J774 macrophages stably knocked down for Coro1A (hereafter referred to as J774 ^Coro1A^ ^KD^) were used (Fig. 6C) and (Fig.S6B) (Jayachandran et al., 2008). The interaction between Coro1A and TGF-βRI was validated in J774 ^WT^ and J774 ^Coro1A^ ^KD^ cells by antibody-based pull-down. The blots revealed interaction of Coro1A and TGF-βRI in J774 ^WT^ cells but not in J774 ^Coro1A^ ^KD^ cells (Fig. 6D).

Under the basal level, there is no significant change in the expression of prototypical M2 markers like Arginase-1, IL-10, master regulator of M2 macrophage markers SNAIL, and TGF-βRI, in J774^WT^ and J774^Coro1A^ ^KD^ cells. Upon stimulation with TGF-β, J774^WT^ exhibited strong induction of M2 markers Arginase-1 (6.877±1.372) and SNAIL (5.395±0.856), whereas this phenotype was severely blunted in J774^Coro1A-KD^ cells (Arginase-1 0.5067±0.0898, SNAIL 2.201±1.164) measured by qRT-PCR (Fig.6E, Left and Right). In line with these changes in gene expression, the secretion of the anti-inflammatory cytokine IL-10 was significantly higher in J774^WT^ cells (55.83±14.29) after TGF-β stimulation; however, in J774 ^Coro1A^ ^KD^ cells treated with TGF-β, IL-10 secretion was considerably lower (6.904±15.23). (Fig. 6F). Furthermore, Arginase-1-positive cells were quantified using Flow Cytometry, revealing that J774 ^WT^ cells showed a pronounced response to TGF-β stimulation, with increased Arginase-1-positive cells (27.67±9.615), whereas J774 ^Coro1A^ ^KD^ cells failed to respond to TGF-β with the same efficiency (4.667±2.749). (Fig. 4G). Considering reduced expression of Arginase, SNAIL, and less secretion of IL-10, the expression of TGF-βRI in J774 ^WT^ and J774 ^Coro1A^ ^KD^ cells was examined. TGF-βRI showed comparable basal levels of expression in both J774 ^WT^ and J774 ^Coro1A^ ^KD^ cells. Upon TGF-β exposure, it was upregulated in both cell types (1.993±0.1658; 1.513±0.4103) (Fig. 6H). This indicates that TGF-β receptor transcription is intact in J774 ^coro1A^ ^KD^ cells, and that impaired signaling likely arises from defects at the protein level. Mechanistically, in the canonical TGF-β signaling pathway, the activated TGF-β receptor phosphorylates SMAD2/3, which form a complex with SMAD4 and translocate to the nucleus to drive transcription of target genes (Nakao & Imamura, 1997). Accordingly, TGF-β stimulation increases the nuclear accumulation of pSMAD2/3 in cells responding to the TGF-β trigger, as compared to less or non-stimulated cells. In line with the above observation, J774 ^WT^ cells exposed to TGF-β showed enhanced nuclear accumulation of SMAD3 as compared to J774 ^Coro1A^ ^KD^ cells (Fig. 6I). Furthermore, bone marrow-derived macrophages (hereafter referred to as BMDMs) isolated from WT mice showed promising accumulation of pSMAD3 in their nucleus on TGF-β stimulation, whereas Coro1A^KO^ mice-derived BMDMs showed impaired nuclear accumulation of pSMAD3, as shown by immunofluorescence microscopy (Fig. 6J). Together, these findings demonstrate that Coro1A is a critical regulator of TGF-β-mediated SMAD3 activation in macrophages and, consequently, of TGF-β-mediated macrophage polarization.

### Coro1A is required for the stability of TGF-βRI

To understand the relevance of Coro1A interaction with TGF-βRI, TGF-βRI expression was investigated in stable J774 ^Coro1A^ ^KD^ cells. A significant decrease in TGF-βRI protein expression levels was seen in J774^Coro1A^ ^KD^ cells compared to J774^WT^ cells (Fig. 7A). Additionally, TGF-βRI expression was validated in BMDM^WT^ and BMDM ^Coro1A^ ^KO^ by confocal microscopy. A significant reduction in FITC-labelled TGF-βRI fluorescence intensity in BMDM ^Coro1A^ ^KO^ was observed, as compared to BMDM ^WT^ (Fig. 7B). Furthermore, to assess overall TGF-βRI-mediated signaling in J774 ^Coro1A^ ^KD^ cells, pAKT/AKT level, as one of the immediate downstream readouts of the TGF-β signaling, was examined. TGF-β-induced pAKT level was markedly reduced in J774 ^Coro1A^ ^KD^ cells as compared to J774 ^WT^ cells, indicating impaired SMAD-independent signaling as well in J774 ^Coro1A^ ^KD^ cells (Fig. 7C). TGF-β signaling is autoregulatory; the activation of intact TGF-β receptors induces *TGFB1* transcription, leading to enhanced precursor synthesis and the release of mature TGF-β into the supernatant. Conversely, impaired TGF-βR signaling lowers both intracellular precursor levels and the secreted TGF-β (Deng et al., 2024; Massagué, 2012).ELISA and immunoblot verified autoregulatory secretion of TGF-β in cell supernatant from J774 ^WT^ and J774 ^Coro1A^ ^KD^ cells. A considerably lower concentration of TGF-β appeared in the cell supernatant collected from J774 ^Coro1A^ ^KD^ cells as compared to J774 ^WT^ cells (Fig. 7D). These findings were further supported by immunoblot analysis of secreted TGF-β in the supernatant, confirming a reduced secretion of TGF-β from J774 ^Coro1A^ ^KD^ cells (Fig.S6C). Further, the TGF-β level in the tissue lysate of WT and Coro1A ^KO^ CAC mice model was measured using ELISA. TGF-β levels were upregulated in CC ^WT^ and CAC ^WT^ compared to both WT Control and their respective Coro1A ^KO^ counterparts, indicating a marked reduction in TGF-β synthesis in Coro1A ^KO^ mice. (Fig. 7E). Next, the mechanism by which Coro1A safeguards TGF-βRI from degradation was investigated. Previous studies have shown that Coro1A decorates signaling endosomes of the neuronal growth factor (NGF)-tropomyosin-related kinase type-1 (TrkA) once it is internalized within the neuronal cell body from lysosomal degradation. (Suo et al., 2014).

**Figure 7.**
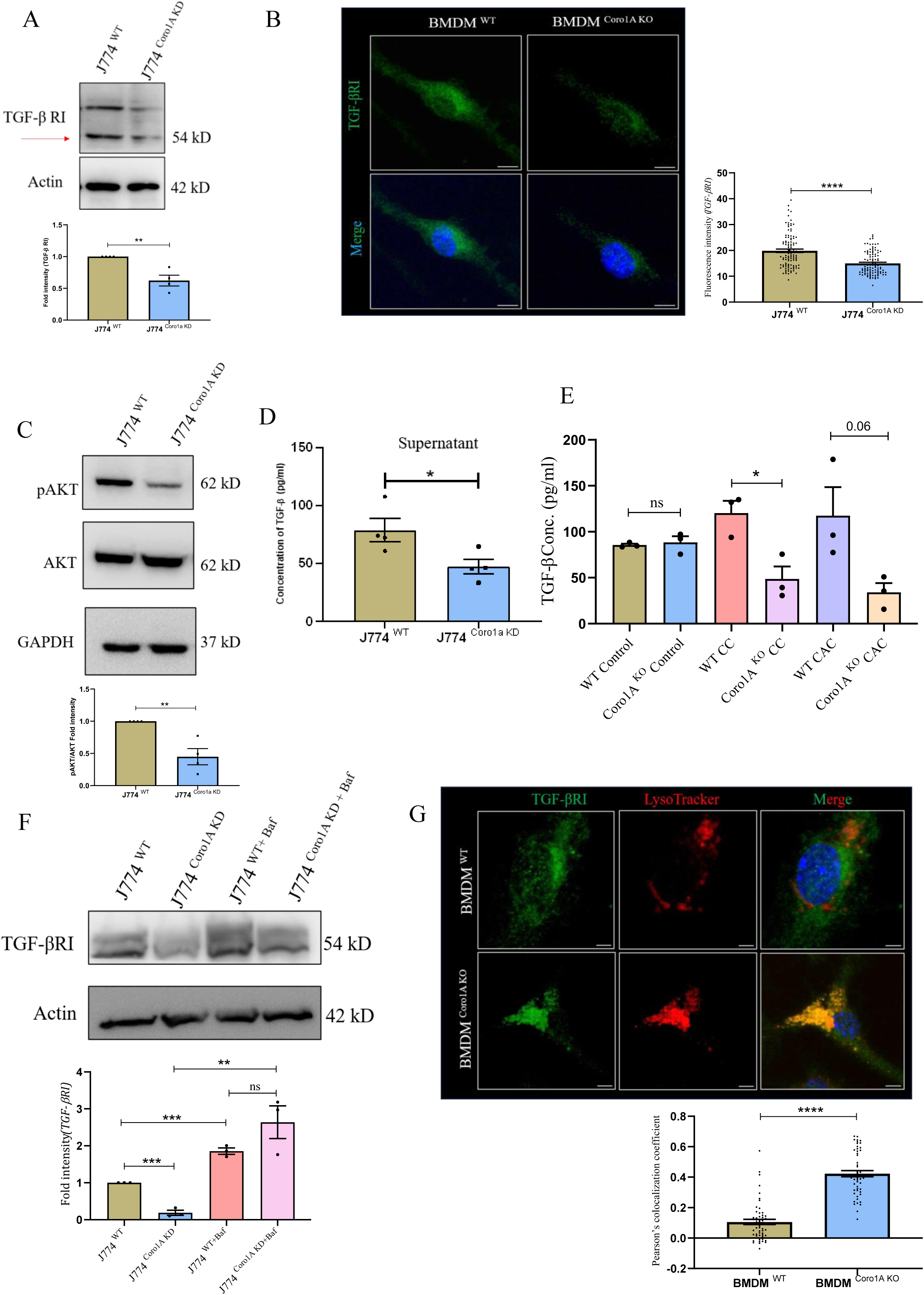
Coro1A is required for the stability of TGF-βRI. A. Immunoblot showing TGF-β R1 expression in J774 ^WT^ and J774 ^Coro1A^ ^KD^ cells. The graph at the bottom shows densitometric analysis of the fold intensity of TGF-βRI, normalized to the Control. B. Representative confocal image of BMDM ^WT^ and BMDM ^Coro1A^ ^KO^ cells stained for TGF-βRI (Green) (n=100). The graph on the right shows the quantification of TGF-βRI mean fluorescence intensity. (Scale bar=5μm). C. Immunoblot showing pAKT and AKT expression in J774 ^WT^ and J774 ^Coro1A^ ^KD^ cells. The graph at the bottom shows the desitometric analysis of the fold intensity of pAKT, normalized to the AKT. D. Concentration analysis of TGF-β in the cell culture supernatant of J774 ^WT^ and J774 ^Coro1A^ ^KD^ cells using ELISA. E. Concentration analysis of TGF-β from the tissue lysates of WT and Coro1A ^KO^ CAC (AOM-DSS) mice model. F. Immunoblot showing TGF-βRI expression in J774^WT^ and J774^Coro1A^ ^KD^ cells with and without Bafilomycin-A1 treatment (100nM). The graph at the bottom shows densitometric analysis of TGF-βRI, normalized to the Control. G. Representative confocal images of BMDM ^WT^ and BMDM ^Coro1A^ ^ko^ cells stained for TGF-βRI (Green) using anti-TGF-βRI antibody and Lysosomes with LysoTracker Red DND-99 (Red). The graph at the bottom illustrates the quantification of colocalization between TGF-βRI and Lysosomes, as determined from images (n=60), using Mander’s overlap coefficient. (Scale bar=5μm). Each dot represents an independent experiment (A, C, D, and F), one cell (B and G), and one mouse (E). Actin was used as a loading control (A and F), and GAPDH was used as a loading control (C). The immunoblots shown here are representative of three biological replicates. Error bars represent mean ± SEM. Statistical analysis performed using the unpaired Student’s t-test, with a p-value <0.05 considered statistically significant. *<0.05, **<0.01, ***< 0.001, ****<0.0001, ns: non-significant.

We hypothesized that the TGF-β-TGF-βRI complex, upon internalization, is protected from degradation by Coro1A in macrophages. In line with this, bafilomycin-A1-mediated blocking of lysosomal acidification rescued the TGF-βRI level in J774 ^Coro1A^ ^KD^ cells as compared to untreated J774 ^Coro1A^ ^KD^ cells (Fig.7F), whereas blocking proteasomal degradation by MG-132 treatment resulted in no change in TGF-βRI expression in J774 ^Coro1A^ ^KD^ cells. (Fig.S6D). As a positive control, p62 accumulated under bafilomycin-A1 treatment, while polyubiquitinated proteins were enriched following MG-132 treatment, confirming effective lysosomal and proteasomal inhibition, respectively (Fig. S6E and F). Furthermore, colocalization of FITC-labelled TGF-βRI(Green) with Lysotracker (Red) in BMDM ^WT^ and BMDM ^Coro1A^ ^KO^ revealed a higher degree of TGF-βRI accumulation within lysosomes of BMDM ^Coro1A^ ^KO^, as observed by strong yellow color puncta (Fig. 7G). Together, these results suggest that in the absence of Coro1A, canonical and non-canonical TGF-β signaling are hampered because TGF-β-TGF-βRI signaling endosomes are predominantly targeted for lysosomal degradation.

### Coro1A utilizes the calcium-calcineurin signaling pathway to protect TGF-βRI from degradation

Previous reports have demonstrated that Coro1A-mediated calcium influx protects NGF-TrkA signaling endosomes and *M. tuberculosis* phagosomes from lysosomal degradation (Jayachandran et al., 2007; Mueller et al., 2008; Suo et al., 2014) (Giorgio Ferrari, 1999).

To test whether Coro1A utilizes calcium ion influx to stabilize TGF-β-TGF-β RI signaling endosomes from lysosomal degradation, RAW 264.7 cells were treated with cell-permeable calcium chelator 1,2-bis(o-aminophenoxy) ethane-N, N, N, N-tetraacetic acid, tetraacetoxymethyl ester (BAPTA-AM) and assessed the TGF-βRI level by immunoblotting. Notably, the level of TGF-βRI in RAW264.7 cells treated with BAPTA-AM was significantly lower than that in untreated controls (Fig. 8A). The result phenocopied the finding observed in the Coro1A-depletion condition. (Fig.7A). Moreover, increasing calcium concentration in J774 ^Coro1A^ ^KD^ cells using calcium ionophore Ionomycin partially rescued TGF-β RI level as compared to J774 ^Coro1A^ ^KD^ cells (Fig. 8B). Taken together, these data suggest stability of TGF-βRI is directly linked to the ability of Coro1A to induce calcium release. Provided that *M*. *tuberculosis* (Jayachandran et al., 2007) and NGF-TrkA signaling endosomes (Suo et al., 2014) utilize calcium ion-activated calcineurin phosphatase to evade lysosomal degradation, it was speculated that TGF-β RI signaling endosomes may employ a similar mechanism. In RAW264.7 cells, potent calcineurin inhibitors, Cyclosporin A (CsA) and FK506, were added to test whether inhibiting calcineurin phosphatase results in degradation of TGF-β RI. Similar to BAPTA-AM, treatment with CsA and FK506 resulted in a significant reduction in TGF-βRI expression (Fig. 8C and D). Collectively, these results suggest that Coro1A, associated with TGF-β-TGF-β RI signaling endosomes, utilizes the calcium-calcineurin signaling pathway to protect TGF-β RI signaling endosomes from lysosomal degradation in the macrophages.

**Figure 8.**
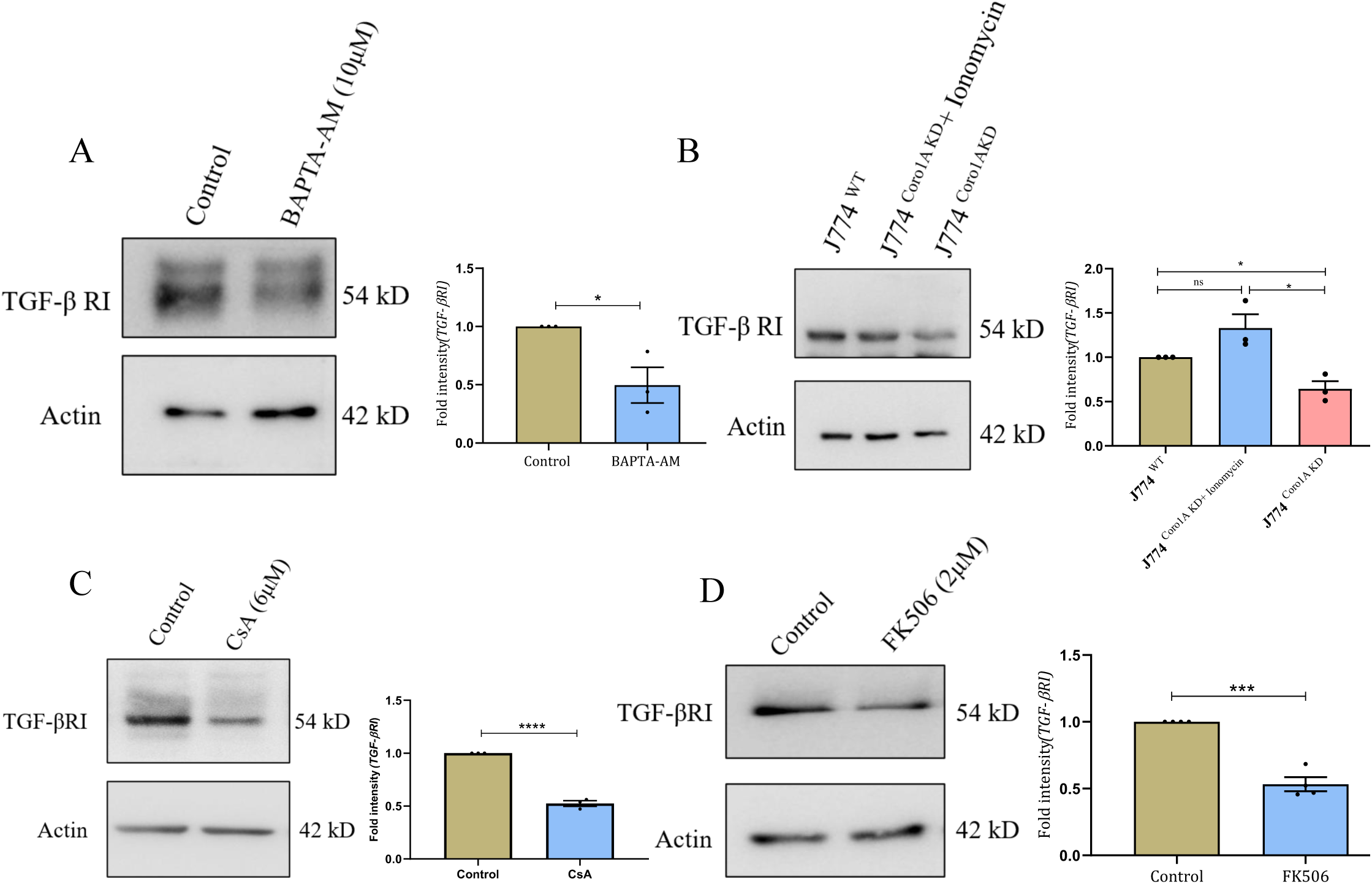
Coro1A utilizes calcium-calcineurin signaling to protect TGF-βRI from lysosomal degradation. A. Immunoblot showing TGF-βRI expression in RAW 264.7 cell lysate after treatment with BAPTA-AM (10 μM). The graph on the right shows densitometric analysis of fold intensity of TGF-βRI, normalized to the loading Control. B. Immunoblot showing TGF-βRI expression in J774 ^WT^ and J774 ^Coro1A^ ^KD^ cell lysates post-treatment with Inomycin (3.5 μM). The graph on the right shows densitometric analysis of fold intensity of TGF-βRI, normalized to the loading Control. C. Immunoblot showing TGF-βRI expression in RAW264.7 cell lysate post-treatment with Cyclosporin A (CsA) (6 μM). The graph on the right shows densitometric analysis of fold intensity of TGF-βRI, normalized to the loading Control. D. Immunoblot showing TGF-βRI expression in RAW 264.7 cell lysate post-treatment with FK506 (3 μM). The graph on the right shows densitometric analysis of fold intensity of TGF-βRI, normalized to the loading Control. Each dot represents an independent biological replicate (A, B, C, and D). As a loading control, Actin was used (A, B, C, and D). The immunoblots shown here are representative of three biological replicates. Error bars represent mean ± SEM. Statistical analysis performed using the unpaired Student’s t-test, with a p-value <0.05 considered statistically significant. *<0.05, **<0.01, ***< 0.001, ****<0.0001, ns: non-significant.

### SUMO2/3 governs the stability of Coro1A

Since Coro1A was identified in this study as an interactor of deSUMOylase SENP5, we tested whether Coro1A is SUMO-modified. Co-immunoprecipitation was performed using anti-SUMO2/3 antibody in J774^WT^ cell lysates, followed by immunoblotting against Coro1A. Interestingly, Coro1A was detected in the SUMO2/3 pull-down (Fig. 9A, bottom). Moreover, the Coro1A band appeared at its usual molecular weight, with no shift in molecular weight detected, indicating that Coro1A itself does not undergo SUMOylation. Surprisingly, in reverse Co-IP with Coro1A, SUMO2/3 bands appeared in the range of 75 kDa-150 kDa, suggesting that Coro1A may interact with already SUMOylated proteins (Fig. 9A, Top). In line with previous findings on Coro2A, where it interacts with SUMOylated Liver X-receptors (LXRs) through SUMO-interacting motifs (SIM)(Huang et al., 2011). It was speculated that Coro1A may utilize its SIM site to interact with SUMOylated proteins. SIM, a conserved hydrophobic amino acid (V/I V/I X V/I), followed by acidic residues, enables the protein to interact with other SUMOylated proteins, influencing their stability, localization, and downstream signaling.(Kerscher, 2007; Song et al., 2004).

**Figure 9.**
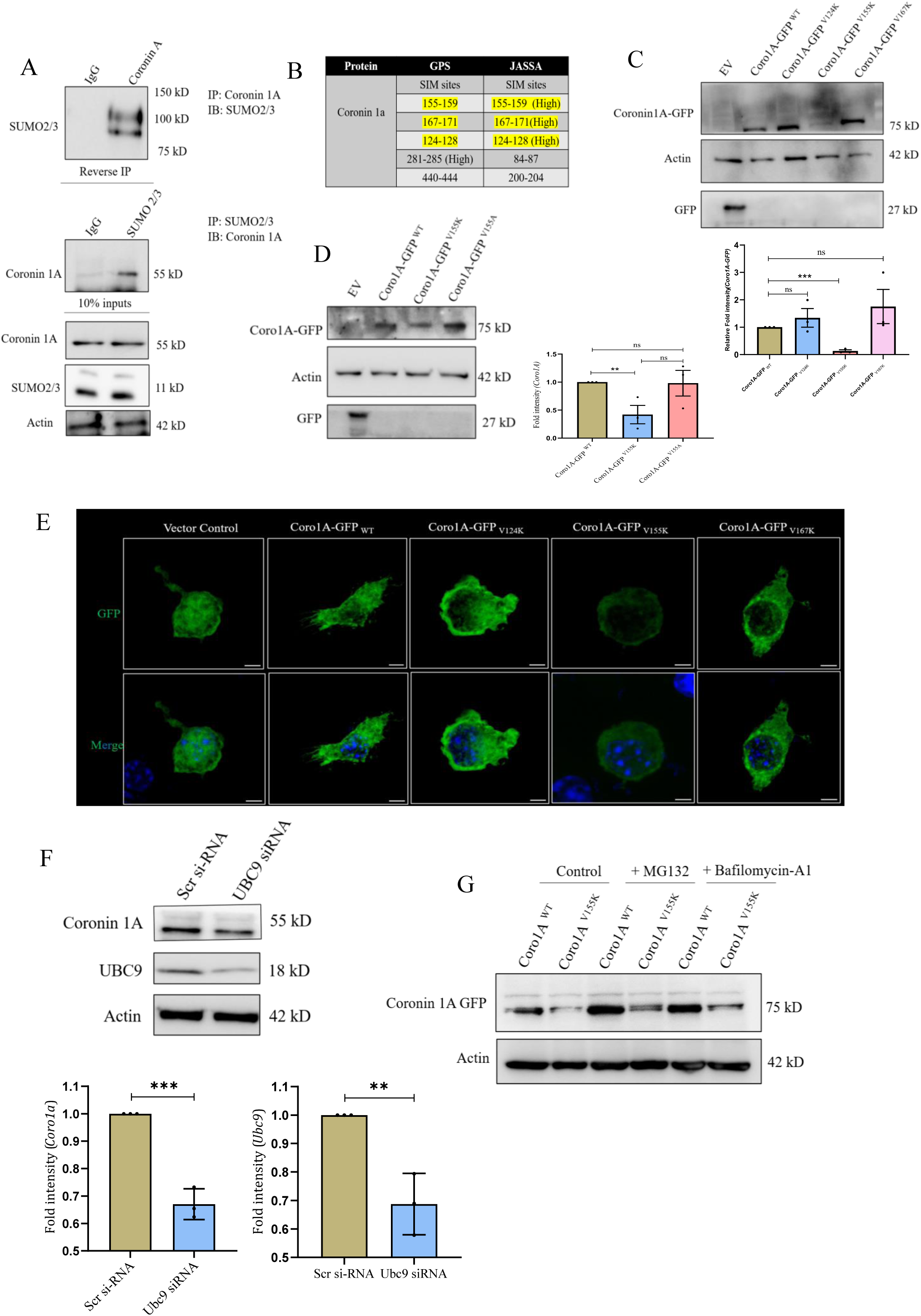
SUMO2/3 governs the stability of Coro1A. A. Top: J774^WT^ cell lysates immunoprecipitated with anti-Coro1A antibody and immunoblotted for SUMO2/3. Bottom: reverse Co-IP using anti-SUMO2/3 antibody from the same lysates and probed for Coro1A with 10% of input loaded and immunoblotted for Coro1A, SUMO2/3, and Actin. Isotype antibodies (IgG) were used as a negative control. B. The table represents possible SIM sites in Coro 1A as predicted by software programs GPS-SUMO and JASSA. SIM sites predicted by both the software are highlighted in yellow. C. Immunoblot showing Coro1A-GFP ^WT^ and SIM mutants (V124K, V155K, V167K) fusion protein expression in transfected RAW264.7 cells. The graph at the bottom shows densitometric analysis of the fold intensity of Coro1A-GFP, normalized to the loading control. D. Immunoblot showing Coro1A -GFP ^WT^ and SIM mutants (V155K, V155A) fusion protein expression in transfected RAW264.7 cells. The graph on the right shows densitometric analysis of the fold intensity of Coro1A-GFP, normalized to the loading control. E. Representative confocal images of Coro1A-GFP ^WT^ and SIM mutants (V124K, V155K, V167K) fusion protein expression in transfected RAW264.7 cells. (Scale bar =5μm). F. Immunoblot showing Coro1A and UBC9 expression upon siRNA-mediated UBC9 knockdown in RAW264.7 cells. The graphs at the bottom show densitometric analysis of the fold intensity of Coro1A and UBC9, normalized to the loading control. G. Immunoblot showing expression of Coro1A-GFP ^WT^ and SIM mutant (V155K) fusion protein expression in RAW264.7 cells with and without treatment of MG-132 (20 μM) and Bafilomycin-A1 (100 nM). Each dot represents an independent experiment (C, D, and F) of biological replicates. Actin was used as a loading control (A, C, D, F, and G). The immunoblots shown here are representative of three biological replicates. Error bars represent mean ± SEM. Statistical analysis performed using the unpaired Student’s t-test, with a p-value <0.05 considered statistically significant. *<0.05, **<0.01, ***< 0.001, ****<0.0001, ns: non-significant.

Using GPS-SUMO and JASSA (Beauclair et al., 2015; Gou et al., 2024),we identified potential SIM sites in Coro1A; however, sites recognized by both tools with high scores were prioritized (Fig.9 B). SIM Sites 124-128, 155-159, and 167-171 were selected for further study. These findings raise two questions: the functional relevance of Coro1A SIM sites and the identity of the protein undergoing SUMOylation to which Coro1A interacts via its SIM site. Conservation of the chosen SIM sites was analyzed using a multiple-sequence alignment across different species (Fig. S7A). To identify the critical SIM site required for Coro1A to function and the relevance of SIM-SUMO interaction, SIM sites were perturbed by performing Site-Directed Mutagenesis (SDM) by replacing the hydrophobic Valine (V) residue with positively charged Lysine (K) residue; this change is expected to disrupt the SIM-SUMO interaction, as established by Song et al. (2004) (Fig.S7 B).

To understand the fate of these mutants, Coro1A-GFP ^WT^ and SIM mutants (Coro1A-GFP ^V124K^, Coro1A-GFP ^V155K^, and Coro1A-GFP ^V167K^) were transfected into RAW264.7 cells, and protein expression was analyzed on an immunoblot. A marked reduction in the expression of Coro1A-GFP^V155K^ was observed on immunoblot (Fig. 9C). To validate the above finding, the hydrophobicity of the SIM site in the Coro1A ^V155K^ mutant was replenished by replacing lysine (K) with alanine (A), resulting in partial rescue of the Coro1A-GFP protein (Fig. 9D). Furthermore, confocal microscopy of RAW264.7 cells transfected with Coro1A-GFP ^WT^ and Coro1A SIM mutants showed that neither mutation altered localization nor induced aggregation in Coro1A. However, Coro1A-GFP ^V155K^ showed a marked reduction in GFP fluorescence intensity as compared to Coro1A-GFP ^WT^ and other SIM mutants (Fig. 9E).

Overall, these results suggest that perturbing SIM sites affects Coro1A stability. Molecular dynamics simulations were carried out between Coro1A ^WT^ and Coro1A ^V155K^ mutant using the CABS-Flex 3.0 server (https://doi.org/10.1093/nar/gkaf412) to rule out the possibility of structural aberration induced by the V155K mutation in Coro1A. The results showed that both the wildtype and V155K mutant proteins exhibited similar dynamic behavior, indicating that the mutation does not induce any significant structural aberration that could cause protein denaturation (Fig. S7C). Next, SIM-SUMO interaction was disrupted independently of Coro1A by transiently knocking down the only SUMO-conjugating enzyme, UBC9, in RAW264.7 cells, thereby lowering global SUMOylation. Under these conditions, Coro1A expression was strikingly less. (Fig.9F). This effect is restricted at the protein level, as the transcript of Coro1A was upregulated on UBC9 knockdown (Fig.S7D). To investigate the route of Coro1A-GFP ^V155K^ degradation, RAW264.7 cells transfected with Coro1A-GFP ^WT^ and ^V155K^ mutant were treated with MG-132 and Bafilomycin-A1. Surprisingly, partial rescue was observed in Coro1A-GFP ^V155K^ in both conditions (Fig.9G). Altogether, these results show that disruption of the SIM site or lowering global SUMOylation leads to degradation of the Coro1A protein, indicating that Coro1A depends on SUMO2/3 and SUMOylated target protein for stability.

### Coro1A interacts with SUMOylated Raftlin-1 through SIM interaction

To identify potential SUMOylated proteins involved in the SIM-SUMO interaction with Coro1A, two distinct bands of SUMO2/3, observed between 75 kD and 150 kD (Fig. 9A, Top), were subjected to mass spectrometry. After subtracting proteins detected in IgG, a list of potential proteins that may undergo SUMOylation was obtained (Fig. 10A).

**Figure 10.**
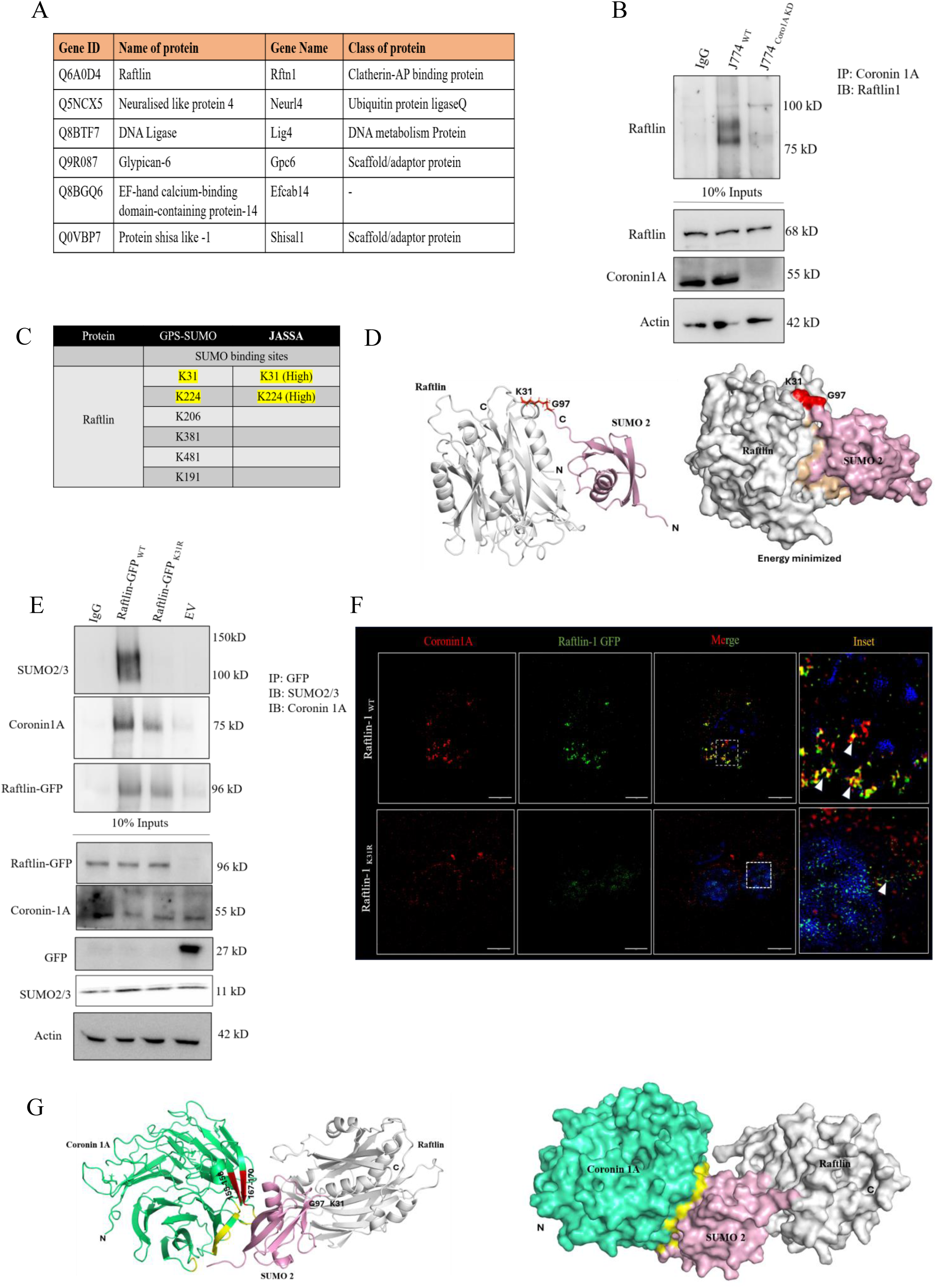
Coro1A interacts with SUMOylated Raftlin-1 through the SIM site. A. The table lists putative SUMOylated proteins identified by mass spectrometry. B. Top: J774^WT^ and J774^Coro1A^ ^KD^ cell lysates immunoprecipitated with anti-Coro1A antibody and immunoblotted for Raftlin. 10% of the input was loaded from the same cell lysate and probed for Raftlin, Coro1A, and Actin. Isotype antibody (IgG) was used as a negative control. Actin was used as a loading control. C. The table shows the possible SUMOylation sites in Raftlin, as predicted by GPS-SUMO and JASSA. SUMoylation sites predicted by both software are highlighted in yellow. D. Computational molecular docking of the alphafold structure of Raftlin and SUMO 2 using HADDOCK2.4. web server. Alpha fold structure of Raftlin (Grey) and SUMO2 is (Magenta). K31, the SUMO binding site is highlighted in Red. E. Top: Raftlin^WT^, Raftlin ^K31R^, and Vector control transfected into RAW264.7 cell lysates were immunoprecipitated with an anti-GFP antibody and immunoblotted for SUMO2/3, Coro1A, and GFP. Bottom: 10% of the input was loaded from the same cell lysate and probed for Raftlin, Coro1A, Actin, and SUMO2/3. Isotype antibody (IgG) was used as a negative control. Actin was used as a loading control. F. Representative images of structured illumination microscopy displaying images of transfected RAW264.7 cells with Raftlin^WT^ and Raftlin ^K31R^ (Green) stained with Coro1A (Red) using anti-Coro1A antibody. The inset shows zoomed areas of images, marked with arrows, to illustrate colocalization (scale bar = 5 μm). G. Computational molecular docking of the crystal structure of Coro1A (2AQ5) and stable binary structure of Raftlin and SUMO 2 performed using HADDOCK2. 4. 3-D structure of Coro1A (Green), Raftlin-1 (Grey), and SUMO2 is Magenta, and SIM sites 155-159 and SIM 167-171 are in red. The area of SUMO2 interacting with Coro1A is highlighted in yellow.

From that list, Raftlin, a novel major lipid raft protein, was selected as a promising candidate to undergo SUMOylation and interact with Coro1A. Previous reports have shown that Raftlin controls T-cell receptor (TCR) and B-cell receptor (BCR) signaling by mobilizing calcium ions (Kazuko Saeki & Akihiko Yoshimura, 2003; Saeki et al., 2009). Raftlin plays a critical role in endocytosis, decorates signaling endosomes, and is detected together with Coro1A in the VEGFR2 interactome (Tatematsu et al., 2016; Watanabe et al., 2011a; Bayliss et al., 2020). These findings indicate a close association between Raftlin and Coro1A, and, together with their functional similarity, led us to select Raftlin as a potential candidate for the Coro1A SIM-SUMO interaction. To test the interaction between Coro1A and Raftlin, co-immunoprecipitation (co-IP) was performed using an anti-Coro1A antibody in J774 ^WT^ and J774 ^Coro1A^ ^KD^ cell lysates. Raftlin-Coro1A interaction appeared in J774 ^WT^, but not in J774 ^Coro1A^ ^KD^ (Fig. 10 B). Furthermore, their interaction was validated in RAW264.7 cells, confirming its conservation across different cell types (Fig. S7E). To identify potential SUMOylation sites present in Raftlin, the GPS-SUMO and JASSA software tools were employed. Both software tools with high scores identified the K31 and K224 sites (Fig. 10C). To narrow down on the possible SUMOylation site, an *in-silico* protein-protein docking approach was employed. While docking the SUMO2 protein on the alpha fold predicted structure of Raftlin, parameters like supramolecular geometry, accessibility of lysine (K) residues for isopeptide bond formation with SUMO, and minimal energy score were considered. Molecular docking demonstrated interaction between the di-glycine motif of SUMO2 (G-97) with K31, K139, and K308 of Raftlin, with SUMO2-K31 interaction being most compact and exhibiting the lowest minimum energy score (Fig. 10D). Additionally, multiple sequence alignment revealed that K31 is highly conserved across different species, underlining its functional importance (Fig.S7F) whereas other mentioned sites are not conserved. To test whether Raftlin undergoes SUMOylation and whether K31 is the potential site of SUMO binding, SDM was performed to substitute lysine (K31) with arginine (K31R), as SUMO can form an isopeptide bond with only lysine and not with arginine (Rytinki & Palvimo, 2009). Next, RAW264.7 cells were transfected with Raftlin ^WT^ and Raftlin ^K31R,^ along with appropriate controls, and subjected to GFP-antibody-based pull-down. Immunoblot revealed a prominent interaction between Raftlin ^WT^ and SUMO2/3, as well as between Raftlin ^WT^ and Coro1A. In contrast, Raftlin ^K31R^ did not interact with SUMO2/3 and showed a feeble interaction with Coro1A (Fig. 10E). Raftlin ^K31R^ showed comparable stability to Raftlin ^WT^ as demonstrated by immunoblotting (Fig. S7G). In line with this, structured illumination microscopy (SIM) demonstrated strong colocalization of Coro1A (Red) with Raftlin ^WT^ (Green), which was markedly reduced with Raftlin ^K31R^ (Fig. 10F). To map the interacting interface of SUMOylated Raftlin with Coro1A, a stable Raftlin-SUMO complex was docked on Coro1A (PDB ID: 2AQ5). Protein-protein docking analysis indicated that SUMO interacts with Coro1A through a region spanning residues 155-159 and 167-171. The SUMO-interacting interface on Coro1A is highlighted in yellow (Fig.10G). Notably, these regions correspond to SUMO-interacting motifs (SIMs) of Coro1A, consistent with our previous findings demonstrating that SIM 155-159 is critical for Coro1A to interact with SUMO2/3 and maintain stability. Overall, these results revealed that Coro1A interacts with SUMOylated Raftlin at K31 through the SIM site.

## Discussion

In the current work, a screen was carried out to assess the possible involvement of SUMOylation machinery components in CAC, leading to the identification of the deSUMOylase- SENP5. To dissect the contribution of SENP5 in CAC, its interactome was analyzed, and Coro1A emerged as an outcome. Coro1A, an actin-binding regulatory protein, is associated with immune cell function but has not been explored in the context of IBD or CAC (Humphries et al., 2002; Mueller et al., 2008; Pick et al., 2017; Stocker et al., 2018). However, increased expression of Coro1A has been reported in several chronically inflamed and autoimmune disease models, such as renal fibrosis, lupus nephritis, and lung fibrosis, as well as in some forms of cancer (X. Deng et al., 2024; Elihamu et al., 2025; Kros et al., 2024; Nicolaou et al., 2020; Niu et al., 2024; Siegmund et al., 2016). In lung adenocarcinoma, increased expression of Coro1A correlated with TNM stage, ranked among the top 10 prognostic markers, particularly as an M2 or TAM macrophage marker, and appeared to shape the tumor microenvironment (TME) (Z. Li et al., 2025; Sun et al., 2024) (Bao et al., 2022; Y. Zhou et al., 2021). Coro1A expression is linked to cancer pathogenesis, since knockdown of Coro1A curbs the growth and metastasis of breast cancer cells (Elihamu et al., 2025; He et al., 2018; Jeon et al., 2024; Kros et al., 2024). Likewise, Coro 1A has been identified as one of the five robust biomarkers in bladder cancer (Kumar et al., n.d.).

A direct interaction of Coro1A with the C-terminus of β2 integrins is required for polymorphonuclear leukocyte (PMNs) trafficking, adhesion, and post-adhesion events (Pick et al., 2017). Furthermore, in a cystic fibrosis model, Coro1A was shown to regulate integrin clustering and activation, a mechanism required for monocyte function (Younis et al., 2024). The immune regulatory function of Coro 1A is exemplified by the finding linking Coro 1A to allograft tolerance. Specifically, T cells lacking Coro1A suppressed the allograft rejection phenotype in a wild-type host upon adoptive transfer, while maintaining T cell-specific responses against the pathogen (Jayachandran et al., 2019). Likewise, Coro1A ^KO^ mice display attenuated response in an experimental model of encephalomyelitis (Jayachandran et al., 2019; Kaminski et al., 2011). Overall, these reports make it evident that several aspects of immune cell function, including maintenance of numbers, trafficking, adhesion, and post-adhesion events, are directly governed by Coro 1AMueller et al., 2008; Ndinyanka Fabrice et al., 2025; Pick et al., 2017; Shiow et al., 2009; Younis et al., 2025) (Siegmund et al., 2016).

Arguably, these phenotypes result from significant signaling alterations in these cells, with Coro1A function playing a direct role. Coro1A was shown to regulate TGF-βR signaling in an experimental autoimmune encephalomyelitis (EAE) model. Coro1A^KO^ mice exhibited less severe EAE. Unexpectedly, upon re-induction of EAE, Coro1A^KO^ mice showed severe disease, with a strong correlation with IL-17-producing T cells. Coro1A^KO^ Th17 cells show a severe defect in TGF-βR-mediated SMAD3 activation, leading to increased IL-17 secretion (Kaminski et al., 2011). TGF-β and IL-4 are key inducers of M2 macrophages and systemic immune suppression. TGF-β promotes immune suppression in T cells, Dendritic cells (DCs), neutrophils, and macrophages via canonical (SMAD2/3-dependent) and non-canonical pathways. TGF-β signaling hampers DC maturation and antigen-presenting capacity by modulating co-stimulatory molecule expression. In T-Cells, TGF-β suppresses TCR-induced cell proliferation and regulatory T cell differentiation by inducing Foxp3. TGF-β also polarizes neutrophils towards the pro-tumorigenic N2 phenotype. Given the broad immunosuppressive effects of TGF-β, we hypothesize that Coro1A may regulate these TGF-β-induced processes in immune cells, as we have demonstrated in macrophages (Z. Deng et al., 2024b; Fridlender et al., 2009; Gong et al., 2012a; C. S. Lin et al., 2013; Mckarns et al., 2004; Strobl & Knapp, 1999).

Owing to TGF-β signaling’s multifaceted functions, mutations in components have also been linked to colorectal cancer, particularly in the tumor microenvironment (TME). Genetic alterations in genes encoding components of the TGF-β signaling pathway, particularly TGF-βRII (50-60%) and SMAD4 (46-48%), are the most common in CAC cases (Calon et al., 2012; Means et al., 2018; Yashiro, 2014). In CAC macrophages, characterized as CD11b^+^ F4/80^+^ Ly6C^+^, are increased and play a crucial role in tumorigenesis; their depletion halts tumorigenesis. Highly plastic macrophages under persistent chronic conditions polarize towards an M2-like phenotype and act as a key driver of the CAC (Hardbower et al., 2017; Sheng et al., 2022; Shin et al., 2023; Wang et al., 2023; Yuan et al., 2021).

Interestingly, polymorphisms in TGF-β Receptors and/or signaling components have been reported to be associated with a concomitant decrease in the CD11b^+^ F4/80^+^ M2 macrophage population (Gong et al., 2012). In line with this, in the current work, we observed that Coro1A is a potent regulator of TGF-β signaling in macrophages in CAC. Myeloid cells play a key role in shaping TME, and TGF-β signaling is essential for myeloid cell function in CAC (Chan et al., 2022). Furthermore, TGF-β-deficient mice exhibit significantly reduced CAC burden, lower tumor burden, and substantially reduced IL-6 and TNF-α production (J. Li et al., 2013; W. Wang et al., 2015). This phenotype closely matches our observation in Coro1A^KO^ mice, which show reduced TGF-β signaling and fewer tumors. The integral role of Coro1A in immune cell function and TGF-β signaling also strongly connects to fibrosis and the mechanisms that contribute to it. The c-AMP-responsive element-binding protein (CREBH) controls Coro1A expression and regulates autophagy in NASH, thereby preventing liver fibrosis. CREBH^KO^ mice exhibit liver fibrosis by activating TGF-β, CTGF signaling pathways, and deposition of collagen, α-SMA, and fibronectin (Deng et al., 2024).An extensive clinical study involving colorectal cancer patients (N=94) showed that overexpression of markers of cancer-associated fibroblasts was associated with tumor-infiltrating lymphocytes, thus revealing a vital role for CAFs in maintaining the colorectal TME (Zadka et al., 2021). In our study, Coro1A^KO^ mice showed resistance to polyp formation in an AOM-DSS model, with significantly reduced markers of fibrosis, accompanied by attenuated proliferation, oncogene, and EMT gene activation. The connection between Coro1A and fibrosis and tumorigenesis was through its regulation of TGF-β signaling. Mechanistically, Coro1A-mediated mobilized calcium activates Calcineurin phosphatases, which, in turn, activate genes involved in preventing lysosomal fusion of phagosomes and signaling endosomes(Jayachandran et al., 2007; Suo et al., 2014). In our study, we also found Coro1A employing the same calcium-calcineurin pathway to protect the TGF-β signaling endosome from lysosomal degradation.

The fascinating part of the work is how SUMOylation of Raftlin protein bridges these diverse pathways. We observed that Coro1A depends on SUMOylated Raftlin for stability via the SUMO-SIM interaction. SIM-deficient Coro1A undergoes rapid degradation, leading to concomitant dysregulated TGF-β signaling. Raftlin and Coro1A exhibit a functional correlation. Like Coro1A, Raftlin also controls B-cell receptor (BCR) and T-cell receptor (TCR) signaling by modulating calcium mobilization, and both are key players in clathrin-mediated endocytosis, a common pathway utilized by TGF-β signaling endosomes to initiate signaling. Given their functional correlation, our finding highlights SUMO as a connecting link between Coro1A and Raftlin (Bayliss et al., 2020; Kazuko Saeki & Akihiko Yoshimura, 2003; Saeki et al., 2009; Tatematsu et al., 2016; Watanabe et al., 2011) (Lacy et al., 2018). Presumably, SENP5 plays a role in Raftlin SUMOylation, an attractive area for future research. Overall, our study revealed a critical role of Coro1A in shaping the tumor microenvironment in CAC by modulating TGF-β signaling and promoting a M2-like non-inflammatory macrophage phenotype.

## Material Methods

### Human patient samples

Human patient samples for IBS (10), UC (19), SCRC (9), and CAC (15) were obtained from the Gastroenterology Department at All India Institute of Medical Sciences, New Delhi, India. Colonic biopsies of patients were obtained during routine colonoscopies, with exclusion and inclusion criteria as specified by Mustafa et al. (2017) and Suhail et al. (2019).Six biopsies were collected from each patient for use in qRT-PCR, immunoblotting, and histopathology. The study included patients with mild and moderate Ulcerative Colitis Endoscopic Index of Severity (UCEIS) scores. Informed consent forms were obtained from each patient during the biopsy procedure. The patient’s sex was not considered in the current study. Ethical approval was obtained from the Regional Centre for Biotechnology (Approval No. RCB-IEC-H-11) and AIIMS (Approval No. IEC-217/05.04.2019, RP-19/2019) for using these patients’ samples in the current study.

### Mice

All C57BL/6 mice were maintained in pathogen-free conditions by following all husbandry standards in the small animal facility of RCB. All mice were provided with sterilized food and water at 25 °C, under a 12-hour light/dark cycle. To induce CAC, Azoxymethane (AOM) was injected intraperitoneally into 6-8-week-old male/ female mice weighing 18-20 g at a dose of 10 mg/kg. To induce chronic colonic inflammation, mice were fed 1% Dextran-sulfate-sodium (DSS) (w/v) (36–50 kDa molecular weight) through autoclaved drinking water for 7 days, followed by a recovery period of 1 week, during which mice were fed normal drinking water. The model lasts for 14 weeks. Mice in the control group were fed with normal drinking water. The same procedure was employed for developing CAC in IL10 ^-/-^ mice (C57BL/6 background, obtained from the National Institute of Immunology, India) and Coro1A^-/-^ mice (C57BL/6 background, kindly provided by Prof. Jean Pieters, Biozentrum, Switzerland). An acute colitis model was performed in 6-8-week-old male/female mice weighing 18-20 g, as described by (Suhail et al., 2019). Symptoms of colitis, such as diarrhea, rectal bleeding, and body weight loss, were monitored routinely throughout the experiment. Sporadic colorectal cancer model performed in APC^+/-^ mice. For 1 week, WT and APC^+/-^ mice were fed 2% DSS, followed by 14 weeks of normal drinking water. Organs, such as the colon and spleen, were examined for gross morphological changes and used for further experimental analysis. For some crucial *in vitro* experiments, Bone marrow-derived macrophages (BMDMs) from WT and Coronin1a ^-/-^ mice were harvested. The RCB Institutional Animal Ethics Committee (IAEC) has approved the above-mentioned animal studies (RCB/IAEC/2024/199, RCB/IAEC/2023/177, RCB/IAEC/2023/156, RCB/IAEC/2021/101).

### Cell culture and transfections

Murine macrophage cell lines RAW 264.7 and J774 ^WT^ and J774 ^Coro1A^ ^KD^ cell lines were grown in DMEM containing 10% fetal bovine serum, 3.7 g/litre Sodium bicarbonate, 1% sodium pyruvate, 1% penicillin and streptomycin solution, and 14 mM HEPES buffer (pH 7.4). BMDMs isolated and cultured from WT and Coronin1A ^-/-^ mice as described by(Toda et al., 2021). All cells were incubated at 37 °C with 5% CO2. RAW264.7 cells were used for plasmid-based overexpression and siRNA-based knockdown studies. WT plasmids and mutants of Coro1A and Raftlin, and Scrambled and targeted UBC9 siRNA were transfected in RAW264.7 using Xtremegene (Merck) and DharmaFECT (Horizon Discovery) according to the manufacturer’s instructions. RAW264.7, BMDMs, J774 ^WT,^ and J774 ^Coro1A^ ^KD^ cells were stimulated with IL-4 (20ng/ml) and TGF-β (10ng/ml) for a specified time period. The pharmacological drugs used in this study to treat RAW264.7, J774 ^WT,^ and J774 ^Coro1A^ ^KD^ cells are MG-132, Bafilomycin-A1, BAPTA-AM, FK-506, Cyclosporin-A, and Ionomycin. Unstimulated or untreated cells served as the control.

### Immunoblot

Human and murine tissues and cell lysates were prepared using RIPA lysis buffer containing 1X protease inhibitor and 20 mM NEM. The BCA assay was used to quantify proteins. Equal concentrations of proteins were separated on sodium-dodecyl sulphate polyacrylamide gel electrophoresis (SDS-PAGE) and transferred to a 0.2 μm or 0.4 μm nitrocellulose membrane. Blots were blocked with 5% skim milk for 1 hour at room temperature, then probed with antibodies against the desired proteins overnight at 4 °C. Protein blots were probed with the following set of antibodies: SENP5, SUMO2/3, SMAD3, Coronin1A, GFP, Actin, GAPDH, TGF-beta, TGF-βRI, pAKT, AKT, Raftlin, Ubiquitin, and P62. The next day, host-specific HRP-conjugated secondary antibodies, anti-Rabbit and anti-Mouse, were added and incubated for 1 hour at room temperature. Protein bands were then detected with Immobilon Forte Western HRP substrate and visualized with the ImageQuant LAS-4000 imaging system. Immunoblot densitometry analysis was performed using ImageJ software.

### Quantitative real-time PCR

RNA was extracted from human patient biopsies, mouse tissue, and different cell lines using the RNeasy RNA isolation kit. cDNA was synthesized using 1 μg of isolated RNA from each sample using the iScript cDNA Synthesis Kit. Real-time PCR was performed using SYBR Green Master Mix in a 20 μL reaction volume on 96-well plates with the CFX96 Real-Time System. The housekeeping genes GAPDH and HPRT were used for normalization based on each sample’s origin. The list of primers used in this study is included in the supplementary material.

### Immunohistochemistry

Mice colon tissue sections were fixed in 4% paraformaldehyde solution overnight and transferred to cryomatrix, followed by snap-freezing in liquid nitrogen. 5 µm-thick sections were cut onto glass slides using a Cryotome. The patient’s archived tissue biopsy slides were obtained from the Department of Pathology at AIIMS, New Delhi, India. Tissue sections were washed three times in 1x PBS (5 minutes each). Further, slides were treated with 3% Hydrogen peroxide for 10 minutes to quench endogenous peroxidase activity, followed by a second wash with PBS. Further slides were blocked with 5% Goat serum for 1 hour. Then, the sections were incubated overnight in a humid chamber with Coronin1A. The next day, sections were washed with PBS, then incubated with HRP-conjugated secondary antibodies for 2 hours. Further slides were washed in PBS, stained with DAB, and counterstained with Hematoxylin. Tissue sections were imaged using a Nikon Bright Field microscope.

### Co-Immunoprecipitation

Tissue lysate and cell lysate prepared using IP lysis buffer having 1X protease inhibitor. Samples were centrifuged at high speed at 4⁰C for 10 minutes to remove debris. The supernatant was quantified to 1 mg protein, followed by the addition of the appropriate antibody at its concentration. Antibody-protein complexes were incubated overnight on an end-to-end rotor. The next day, activated Agarose-A-beads were used to pull down the antibody-protein complex. Beads were sedimented by centrifugation (0.5 rcf, 4⁰C), washed with IP lysis buffer, and then denatured. An isotype control IgG was used as a negative control.

### Nuclear fraction isolation

Cells were washed with chilled PBS, scraped, and collected in pre-chilled tubes on ice. After centrifugation (10,000 rpm, 5 minutes, and 4 °C), the pellet was resuspended in a lysis buffer containing chilled PBS, protease inhibitors (PI), and NP-40, and incubated on ice for 10 minutes. To separate the cytoplasmic fraction, the cell lysate was centrifuged (10 minutes, 10,000 rpm, 4 °C). The supernatant is the cytoplasmic fraction, and the pellet is the nuclear fraction. The obtained nuclear pellet was washed 3 times with lysis buffer and subsequently denatured.

### Hematoxylin and Eosin (H&E) and Sirius red staining

5-μm-thick paraffin-embedded tissue sections were deparaffinized in Xylene and rehydrated through a graded ethanol to distilled water. For H & E staining, tissue sections were stained with Hematoxylin, followed by eosin counterstaining. For Sirius red staining, tissue sections were stained with Weigert’s iron hematoxylin, followed by Sirius red stain, and subsequently washed with acidified water to visualize collagen. Sliders were dehydrated in graded alcohol, cleared in Xylene, and mounted using DPX mountant. Tissue sections were visualized under a bright-field microscope.

### Immunocytochemistry and structured illumination microscopy

Cells seeded on a glass coverslip, fixed with 4% methanol-free paraformaldehyde, washed with PBS, and blocked in 0.1 % BSA containing 0.01% Triton X-100 for 1 hour. Further cells were incubated overnight at 4⁰C with primary antibodies against Coro1A, TGF-βRI, GFP, pSMAD3, Lysotracker, and Phalloidin-594, followed by a fluorochrome-conjugated secondary antibody the next day for 2 hours at room temperature. DAPI was used to counterstain nuclei. Coverslips were mounted using Prolong Gold antifade reagent and imaged by confocal microscopy (Leica SP8) or structured illumination microscopy (Elyra PS1, Carl Zeiss).

### Lamina propria isolation

The Lamina propria of the colon was isolated using the method described by Weigmann et al. (2007). Briefly, the harvested mouse colon was washed with ice-cold PBS, then longitudinally cut open and again washed with PBS to remove mucus. The open intestine is cut into small pieces and subjected to predigestion in a buffer containing 5 mM EDTA and 1 mM DTT for 15-20 minutes in a shaking incubator. With vigorous vortexing, the predigested tissue was filtered through a 100 μm filter. The remaining tissue was finely chopped and incubated in a buffer containing collagen and DNase I (50mg per 100 ml PBS) for one hour at 37⁰C on slow shaking. After incubation, the tubes were vortexed, and the cell suspension was passed through a 40 μm strainer. The collected single-cell suspension was vortexed and centrifuged at 500 g for 10 minutes at room temperature. The pellet obtained was resuspended in PBS, and proceeded to flow cytometric immune phenotyping.

### Flow Cytometry

Flow cytometric analysis was carried out for immune cells retrieved from the lamina propria region of the colon (*in vivo*) and cultured J774 ^WT^ and J774 ^Coro1A^ ^KD^ cells (*in vitro*). *In vivo*-derived immune cells were stained with fixable live-dead stain (Aqua Zombie) for 15 minutes at room temperature in the dark, followed by surface marker staining with fluorophore-conjugated antibodies against CD-45, CD11b, F4/80, and Ly6G for 1 hour at 4⁰C. Further, cells were fixed with 4% paraformaldehyde at room temperature, followed by permeabilization with chilled methanol for 20 minutes on ice. Intracellular staining was performed using anti-iNOS and anti-Arginase-1 fluorophore-conjugated antibodies, along with a primary antibody against Coronin1A for 1 hour of incubation, followed by secondary antibody staining for 1 hour. The same fixation, permeabilization, and intracellular staining procedure was employed for *in vitro*-cultured J774 ^WT^ and J774 ^Coro1A^ ^KD^ cells to stain Arginase-1. Stained cells were resuspended in FACS buffer, acquired on a BD FACSymphony, and analyzed using FlowJo software.

### Enzyme-linked immunosorbent assay

Cell supernatant collected from the cell culture system and colonic tissue lysates were centrifuged to remove debris. The samples were quantified using the BCA assay, and equal concentrations of samples were subjected to a kit-based enzyme-linked immunosorbent assay (ELISA) (R&D Systems) to measure TGF-β & IL-10.

### Protein Digestion and LC-MS/MS

Murine tissue lysates were immunoprecipitated using an anti-SENP5 antibody, and J774 cell lysates were immunoprecipitated with an anti-Coronin1A antibody. An isotype-matched IgG antibody was used as a negative control. The co-immunoprecipitated samples were resolved on 10% SDS–PAGE, followed by in-gel digestion with Trypsin Gold at a 1:50 enzyme-to-substrate ratio. The resulting peptides were extracted and purified using C18 StepPak columns, desalted, and reconstituted in a solution of 98% water, 1% acetonitrile, and 0.1% formic acid.

Tandem MS/MS analysis was performed on a TripleTOF 5600 mass spectrometer (AB Sciex) operated in information-dependent acquisition (IDA) mode. The instrument was coupled to a trap column (Magic C18 AQ, 0.1 × 20 mm, 3 μm, 200 Å) and an analytical column (Magic C18 AQ, 0.1 × 150 mm, 3 μm, 200 Å) via a nano-electrospray ionization source. Peptide separation was achieved using a linear gradient of 5–75% solvent B (100% acetonitrile with 0.1% formic acid) over 135 minutes at a flow rate of 300 nL/min; solvent A was 100% water with 0.1% formic acid.

The mass spectrometer was operated in data-dependent acquisition mode, automatically switching between MS and MS/MS scans. Survey MS spectra were acquired over an m/z range of 300–2000 at a resolution of 75,000. The six most abundant precursor ions per cycle were selected for fragmentation by collision-induced dissociation (CID), with a fixed cycle time of 3 s and a dynamic exclusion period of 2 min. The obtained raw files from the MS system (.wiff) were processed for peptide/protein identification using MaxQuant software (v1.6.0.16) under default settings. An in-house combined UniProt (*Mus musculus*) database, together with common contaminants sequences, was provided for MS/MS spectra search. The detailed workflow of peptide/protein identification, including other search parameters and filtering criteria, was followed as mentioned in (Suhail et al., 2019)

### Structural modeling and docking

The crystal structure of Coro1A (PDB ID: 2AQ5) and the AlphaFold-predicted structures of Raftlin1 and SUMO2 were used for protein–protein docking. Blind docking was performed using the HADDOCK 2.4 web server (https://rascar.science.uu.nl/haddock2.4/) in standard-precision mode, generating multiple poses without predefined binding sites. The docked models were clustered and ranked using the default HADDOCK scoring function, and the top three complexes were selected for refinement. Further, using the Schrödinger software suite (https://www.schrodinger.com/), the energies of poses were minimized, and binding energies were calculated. The final model was selected based on the high docking score and favorable minimized binding energies.

### Multiple sequence alignment (MSA)

FASTA sequences of Coro1A and Raftlin from 8 organisms were retrieved from the Uniprot database and aligned using Clustal Omega with default parameters. Multiple sequence alignments were visualized and analyzed using Jalview (v2.11.5.0) with the ClustalX coloring scheme. Alignment annotations, including consensus, conservation, quality scores, and occupancy, were displayed below the alignment.

### Molecular Dynamics Simulation

The wild-type Coro1A structure (PDB ID: 2AQ5) and the PyMOL-generated Coro1A V155K mutant were subjected to molecular dynamics simulations using the CABS flex server with default parameters for 10 seconds, an equivalent simulation time. The server uses coarse-grained Monte Carlo simulations to efficiently sample near-native protein conformational dynamics. Default parameters were used, including structural clustering and all-atom reconstruction. The resulting ensembles were analyzed for conformational flexibility and residue fluctuations, with residue-wise mobility quantified using root-mean-square fluctuation (RMSF) plots generated with the server’s analysis tools.

### Quantification and statistical analysis

Data were analyzed and visualized using GraphPad Prism version 8.0.1. Results are presented as the mean ± standard error from independent experiments performed in triplicate. As appropriate, statistical analysis was carried out using the unpaired two-tailed Student’s t-test or Welch’s t-test. A *p*-value less than 0.05 was considered statistically significant.

## Acknowledgment

We thank the sophisticated instrumentation facility at the Central Instrumentation Facility (CIF) of the Regional Centre for Biotechnology (RCB), the Flow Cytometry, Confocal Microscopy, Genomics, and Mass Spectrometry facility of the Advanced Technology Platform Centre (ATPC) in the RCB campus, and the Experimental Animal Facility (EAF) at NCR-Biotech Science Cluster for animal experiments. We acknowledge Addgene for the Coro1A plasmid (102815). We are grateful to Prof. Josef M. Penninger and Prof. Jean Pieters for sharing Coro1A knockout mice and J774 ^WT^ and J774 ^Coro1A^ ^KD^ cells. We are thankful to Prof. Venuprasad Poojary for the Raftlin Plasmid. We thank Prof. Tushar K. Maiti for his extended help with mass spectrometry-based experiments. We are grateful to Dr. Rajender Motiani and Dr. Pallavi Keshtrapal for sharing the crucial reagents used in the experiments.

## Author Contributions

**Conceptualization**: Chittur V. Srikanth.

**Methodology**: Rohan Babar, Chittur V. Srikanth.

**Investigation**: Rohan Babar, Poorvi Saini, Neha Guliya, Prakhar Varshney, Vishnu Ashok Kumar, Prabhakar Mujagond.

**Resource**: Rohan Babar, Aamir Suhail, Mukesh Singh, Dolly Jain, Lalita Mehra, Shivangi Tyagi.

**Supervision**: Avinash Bajaj, Krishnan Vengadesan, Prasenjit Das, Vineet Ahuja, and Chittur V. Srikanth.

**Project Administration**: Chittur V. Srikanth.

**Writing-original draft**: Rohan Babar, Poorvi Saini, Neha Guliya, and Prakhar Varshney.

**Writing -review and editing**- Rohan Babar and Chittur V. Srikanth.

---

**Figure 11.**
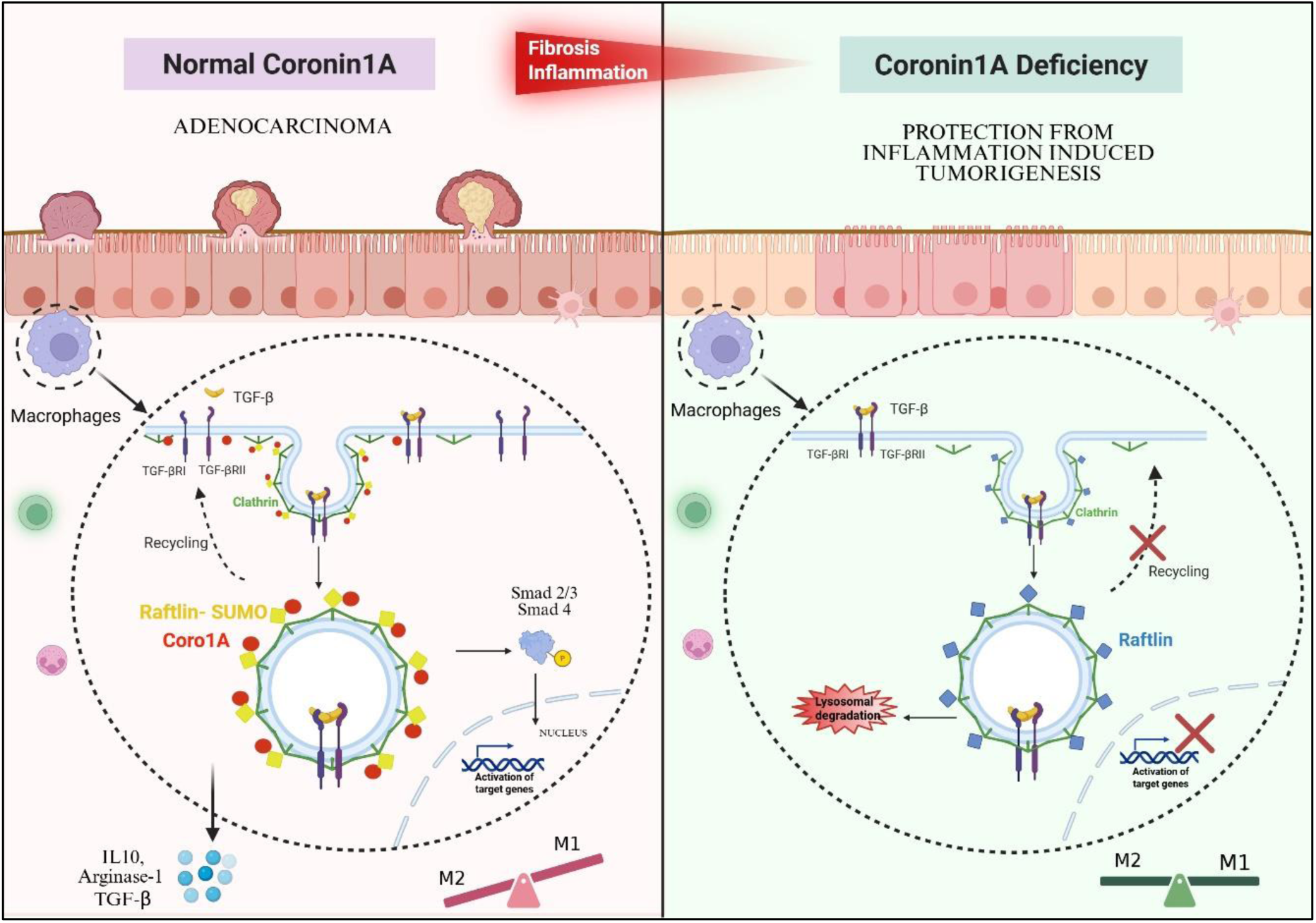
Graphical Summary demonstrating the role of Coro1A in regulating TGF-β signaling and modulating tumor microenvironment.

**Supplementary Figure 1.**
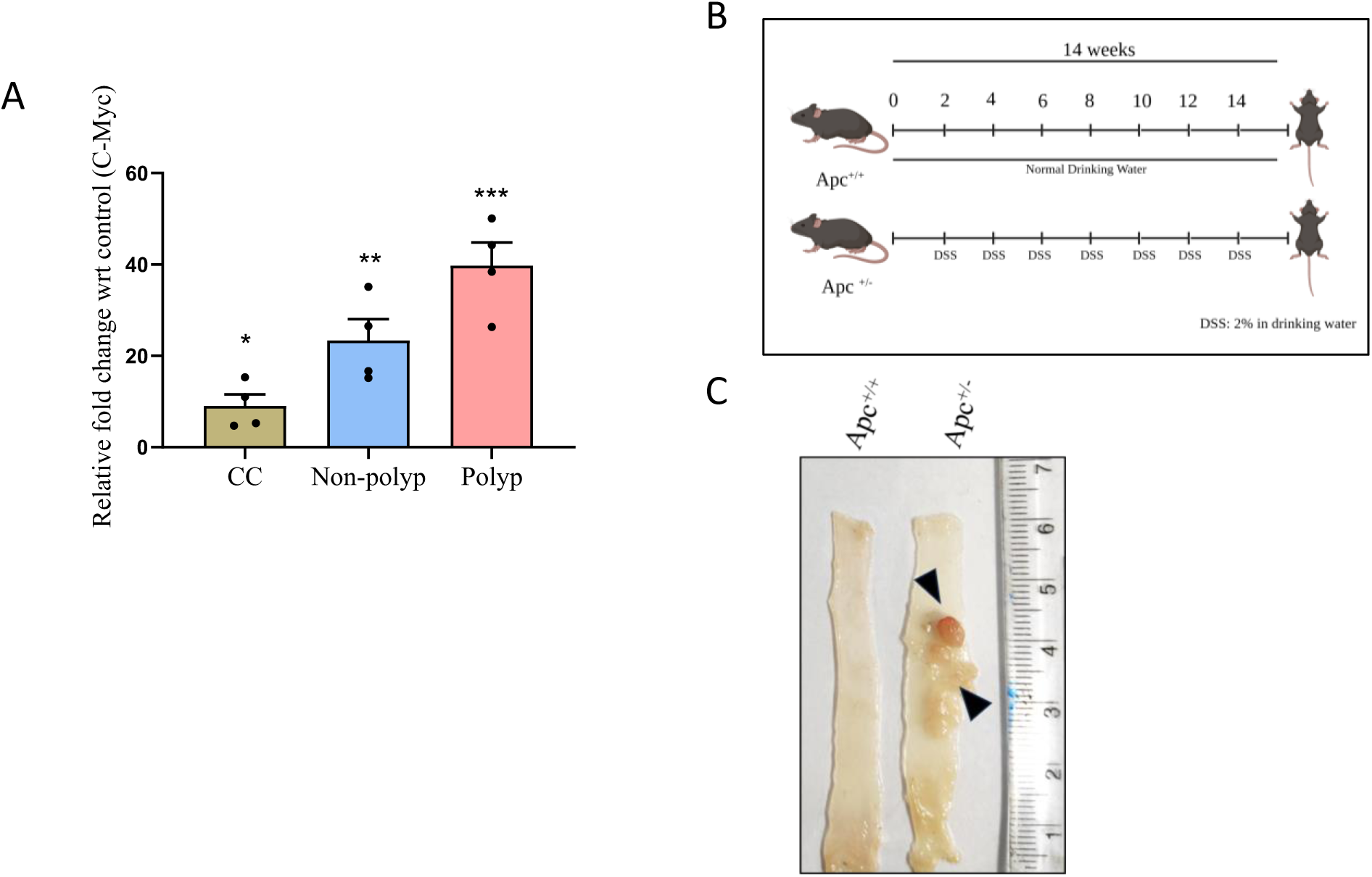
A. qRT-PCR-based fold change expression of C-Myc from CC, non-polyp, and polyp tissue of AOM-DSS mice model. GAPDH was used for gene normalization. Each dot represents one mouse. B. Schematic representation of the sporadic colorectal cancer (SCRC) model in Apc^+/+^ and Apc ^+/-^ mice. C. Representative longitudinal cut-open colon image of Apc^+/+^ and Apc^+/-^ mice. Black arrows indicate polyps present at the distal end of the colon. Error bars represent mean ±SEM. Statistical analysis performed using the unpaired Student’s t test, with a p-value <0.05 considered statistically significant. *<0.05, **<0.01, ***< 0.001, ****<0.0001, ns: non-significant.

**Supplementary Figure 2.**
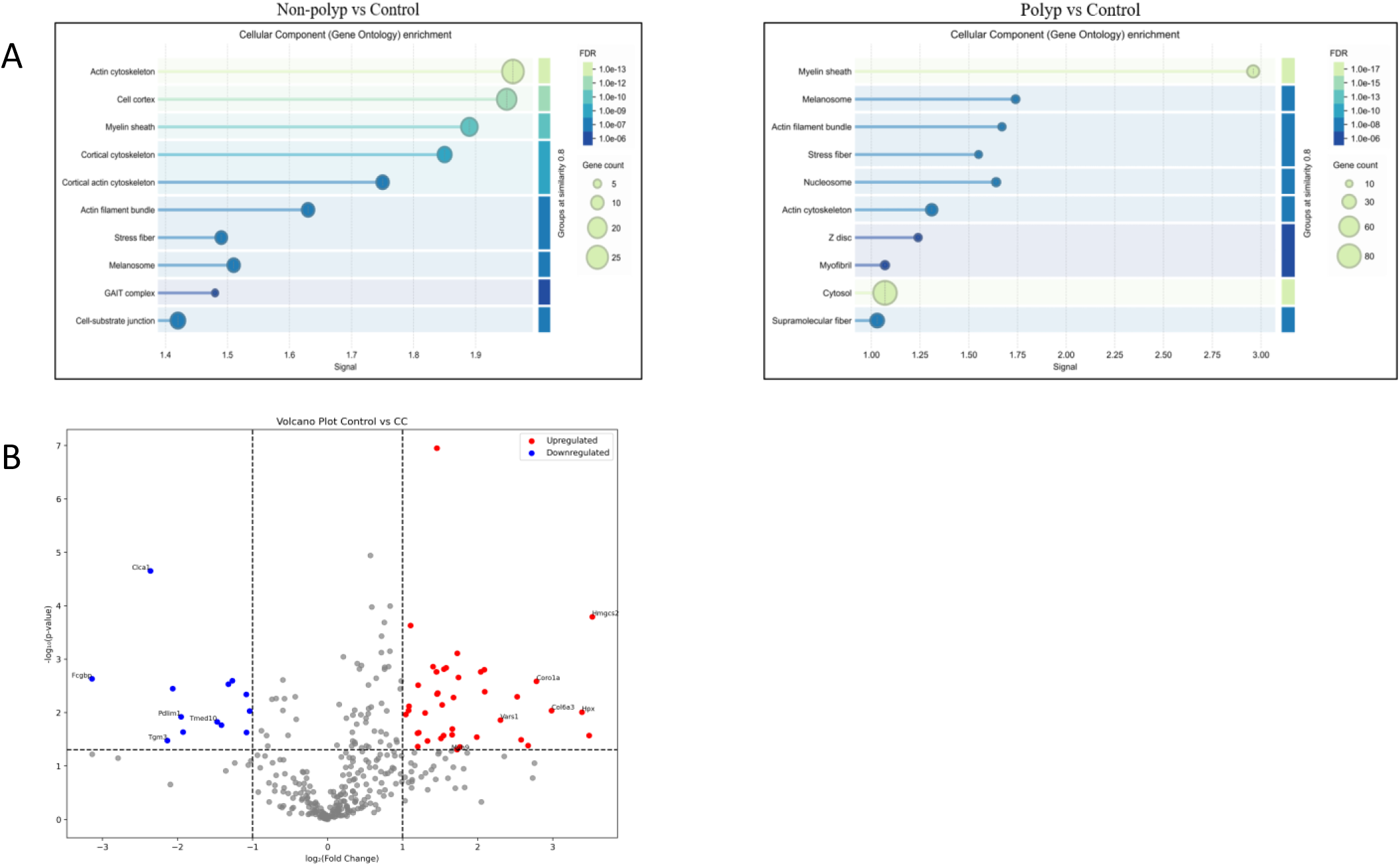
A. The Gene Ontology (GO) Pathway analysis (cellular component) of differentially enriched SENP5 interacting proteins in the non-polyp and polyp region, as compared to the Control. Pathway enrichment significance was assessed using hypergeometric and multiple test correction at FDR less than 0.05. B. Volcano plot of differentially enriched proteins with SENP5 in CC as compared to the Control, with shortlisted potential SENP5 interactors marked. Here, significance is defined as fold change >2 and *p*-value < 0.05.

**Supplementary Figure 3.**
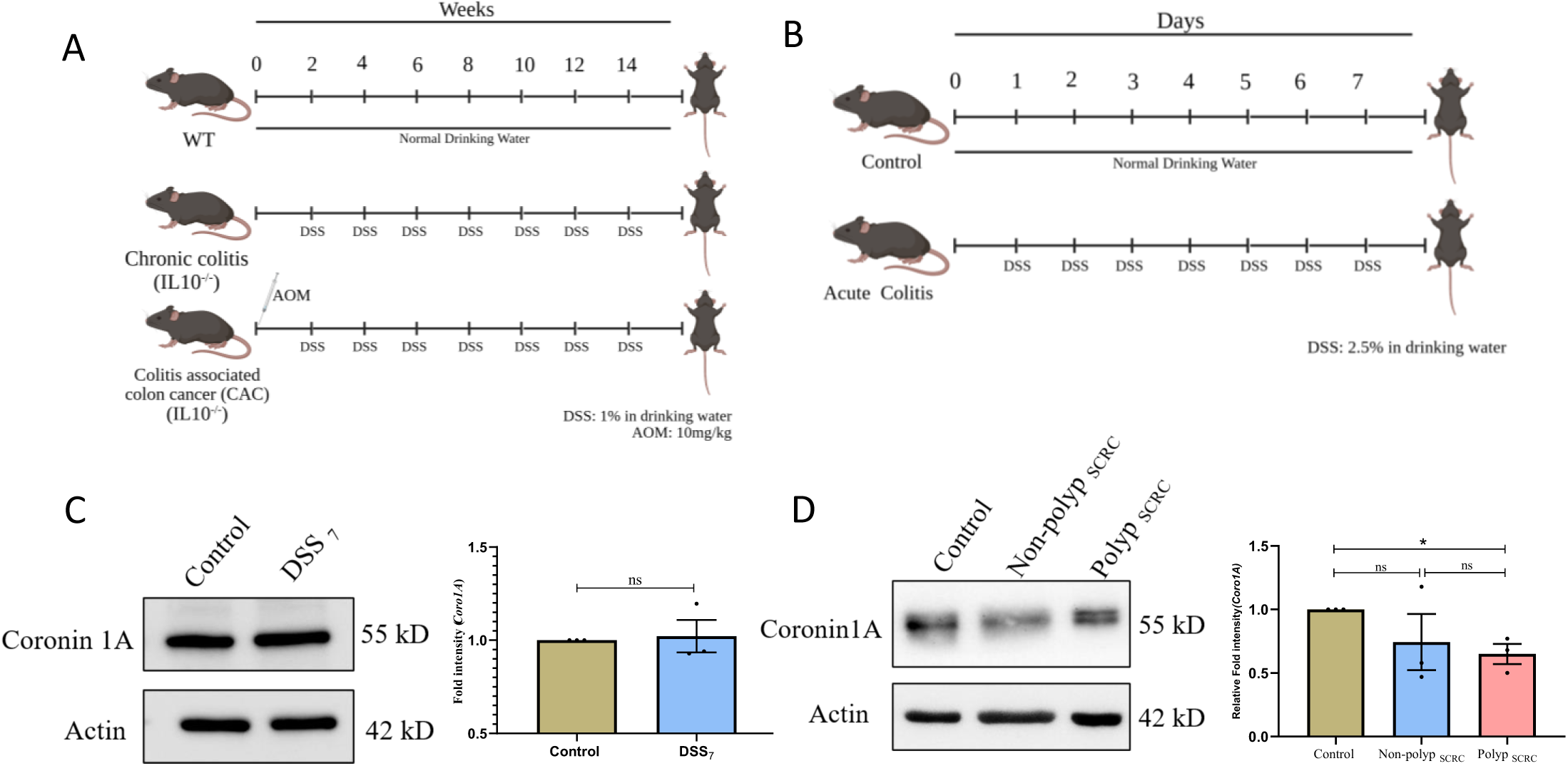
A. Schematic representation of the CAC (AOM-DSS) model in WT and IL10^-/-^ mice. B. Schematic representation of the Acute colitis model (Day-7) in WT mice. C. Immunoblot showing Coro1A expression in lysates of colonic tissue from the acute colitis model. The graph on the right shows densitometric analysis of fold intensity of Coro1A expression, normalized to loading control. D. Immunoblot showing Coro1A expression in colonic tissue lysates of Control, non-polyp, and polyp (Apc^+/-^)SCRC model. The graph on the right shows densitometric analysis of fold intensity of Coro1A expression, normalized to loading control. Each dot represents one mouse (C and D). Actin was used as a loading control (C and D). The immunoblots shown here are representative of three biological replicates. Error bars represent mean ±SEM. Statistical analysis performed using the unpaired Student’s t-test, with a p-value <0.05 considered statistically significant. *<0.05, **<0.01, ***< 0.001, ****<0.0001, ns: non-significant.

**Supplementary Figure 4.**
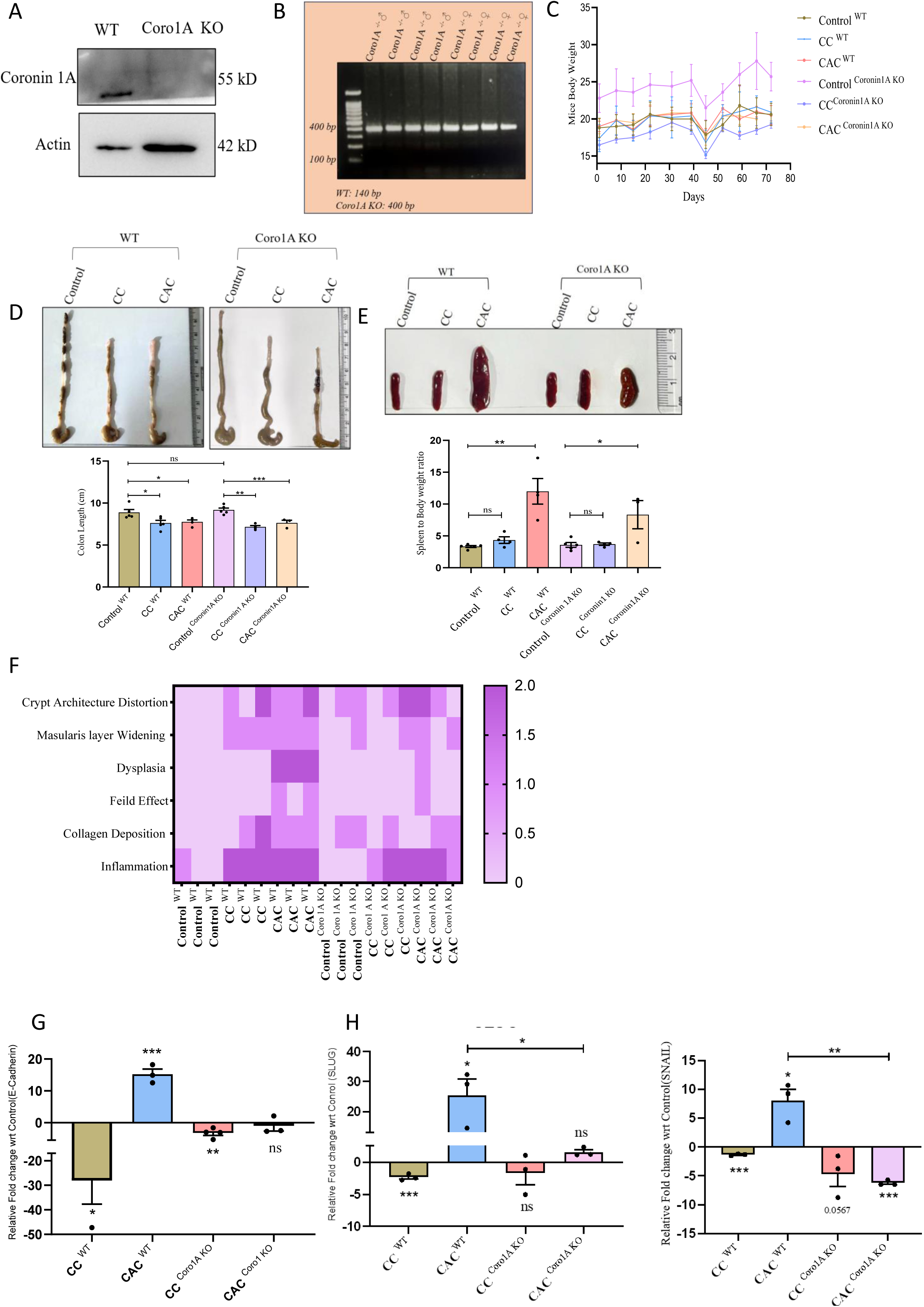
A. Representative immunoblot confirming loss of Coro1A protein expression in colonic tissue lysate of Coro1A ^KO^ mice compared to WT mice. B. Representative agarose gel image confirming the genotype of Coro1A ^KO^ mice (400bp). C. The graph represents the body weight of experimental animals during the development of the AOM-DSS model. D. A representative image showing the gross morphology of the colon from both genotypes (WT and Coro1A ^KO^). The graph at the bottom shows the quantification of colon length. E. A representative image showing the gross morphology of the spleen (Splenomegaly) from both genotypes (WT and Coro1A ^KO^). The graph at the bottom shows the quantification of spleen weight to the mice’s body weight. F. Histopathological scoring of murine colon tissue sections was performed in a blinded manner by an experienced pathologist. Parameters such as collagen deposition, mascularis layer widening, crypt architecture distortion, and inflammation were scored as 0 (absent), 1(mild), and 2 (moderate). Field effect was scored as 0 (absent) and 1 (present), and dysplasia as 0 (absent), 1 (low-grade dysplasia), and 2 (high-grade dysplasia). G. qRT-PCR-based fold change expression of E-Cadherin relative to the average control values. H. qRT-PCR-based fold change expression of EMT markers, SLUG, and SNAIL relative to the average control values. Each dot represents one mouse (D, E, F, and G). Samples analyzed by qRT-PCR include CC ^WT^, CAC ^WT^, CC ^Coro1A^ ^KO^, and CAC ^Coro1A^ ^KO^. GAPDH was used for Gene normalization (F and G). Error bars represent mean ± SEM. Statistical analysis performed using the unpaired Student’s t-test, with a p-value <0.05 considered statistically significant. *<0.05, **<0.01, ***< 0.001, ****<0.0001, ns: non-significant.

**Supplementary Figure 5.**
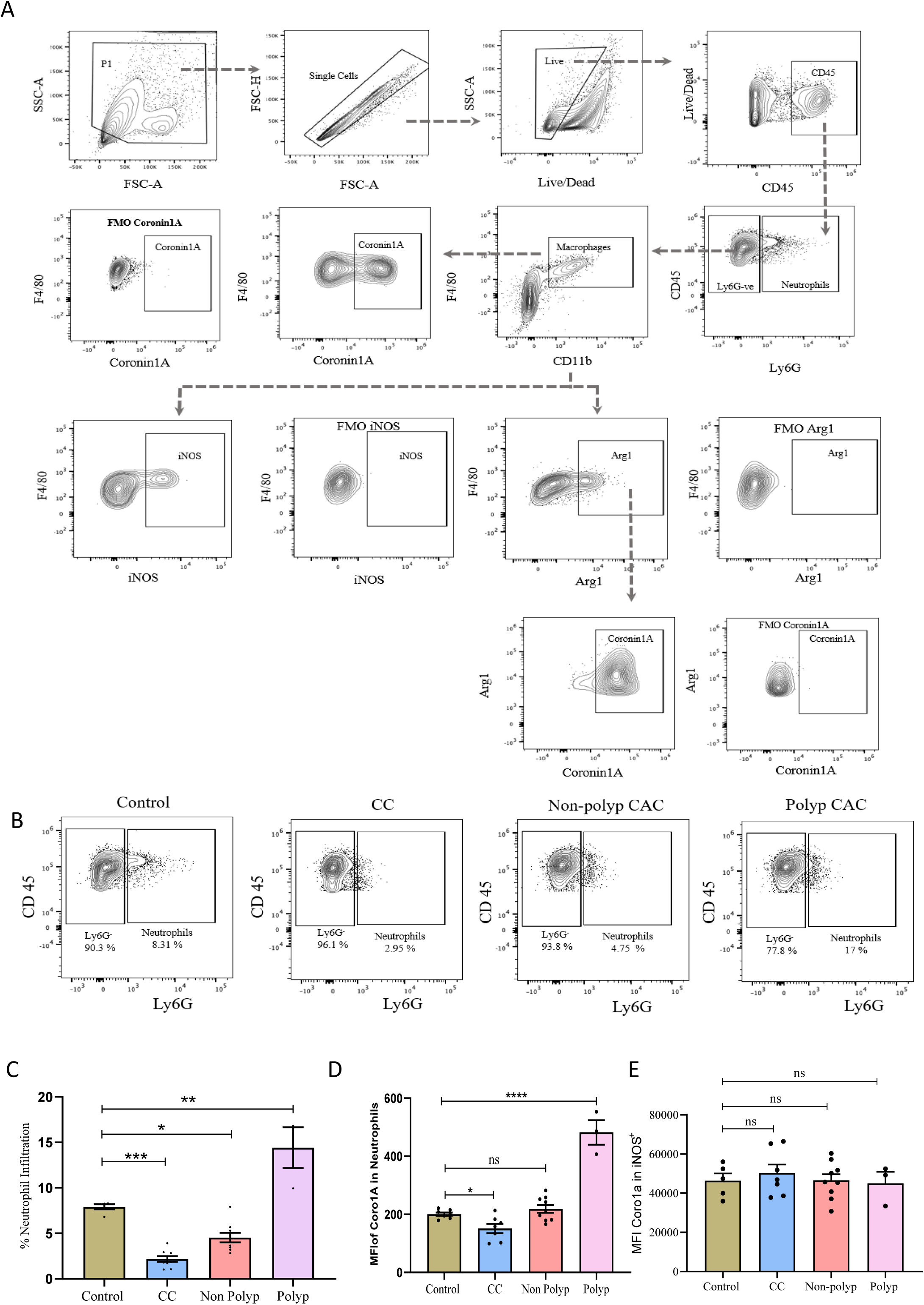
A. Representative flow cytometry plots showing the gating strategy used for cell population identification and analysis, including a fluorescence-minus-one (FMO) control. B. Representative counter plots showing CD 45^+^ and Ly6G^+^ neutrophil infiltration in Control, CC, non-polyp, and polyp regions. C. Percentage infiltration of Neutrophils in Control, CC, non-polyp, and polyp as determined by flow cytometry. D. Median Fluorescence intensity (MFI) of Coro1A in infiltrated neutrophils across Control, CC, non-polyp, and polyp as measured by flow cytometry. E. MFI of Coro1A in iNOS^+^ macrophages across Control, CC, non-polyp, and polyp. Here, each dot represents one mouse (B, C, D, and E). Error bars represent mean ± SEM. Statistical analysis performed using the unpaired Student’s t-test, with a p-value <0.05 considered statistically significant. *<0.05, **<0.01, ***< 0.001, ****<0.0001, ns: non-significant.

**Supplementary Figure 6.**
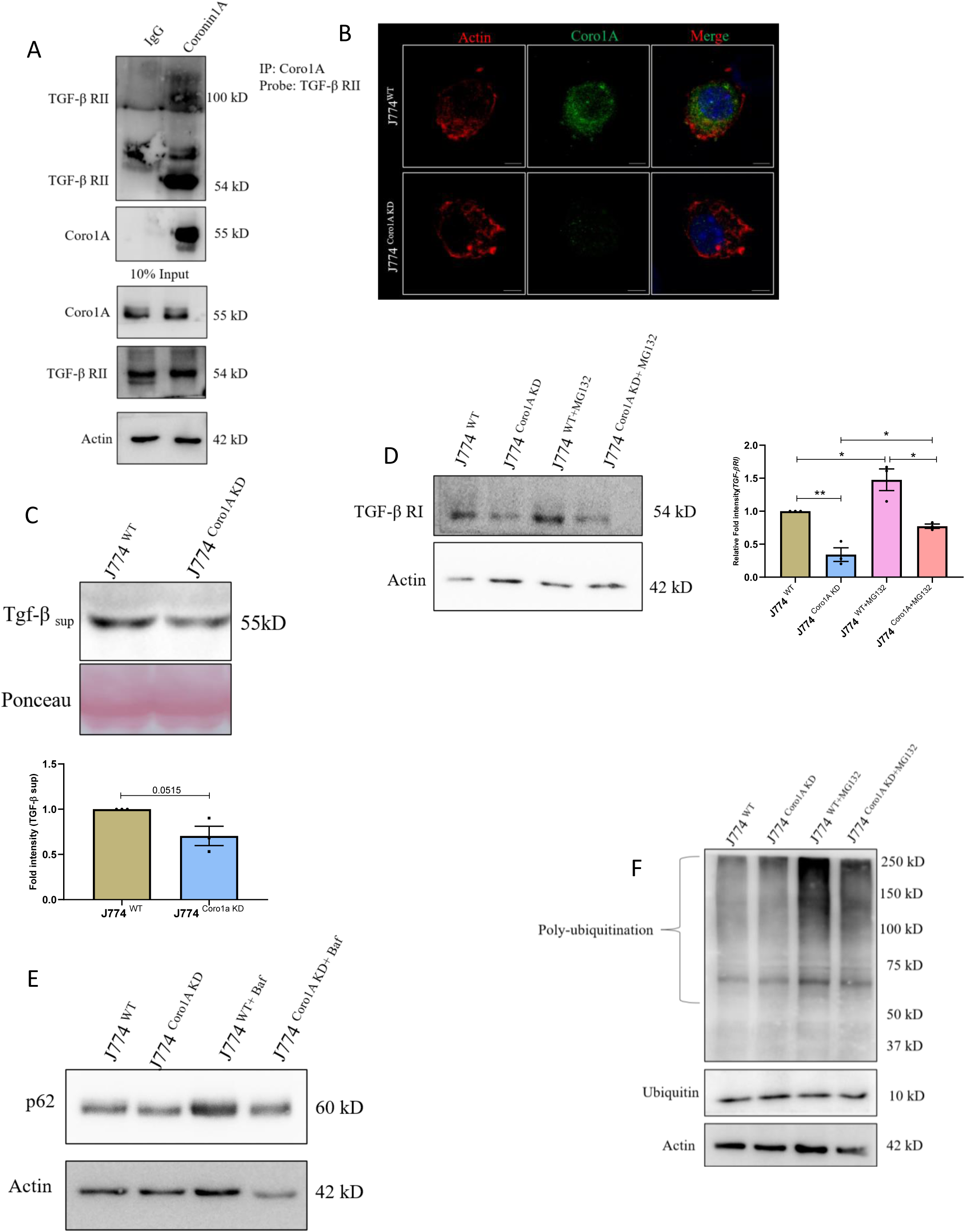
A. Top: RAW264.7 cell lysates immunoprecipitated with anti-Coro1A antibody and immunoblotted for TGF-βRII. Bottom:10% of input loaded from the same cell lysate and immunoblotted for Coro1A, TGF-β RII, and Actin. Isotype antibodies (IgG) were used as a negative control. B. Representative confocal images of J774 ^WT^ and J774 ^Coro1A^ ^KD^ cells showing a marked reduction in the expression of Coro1A in J774^Coro1A^ ^KD^ cells (Scale bar=5μm). Coro1a shows typical cortex-based localization in J774^WT^ cells, which is absent in J774^Coro1A^ ^KD^ cells. Actin was stained with Phalloidin to mark the cell boundaries. C. Immunoblot showing TGF-β secreted by J774 ^WT^ and J774 ^Coro1A^ ^KD^ cells in respective cell culture media. Ponceau image was used as a loading control. The graph at the bottom shows densitometric analysis of fold intensity of Coro1A expression, normalized to the loading control. D. Immunoblot showing TGF-β RI expression in J774 ^WT^ and J774 ^Coro1A^ ^KD^ cells with and without MG-132 treatment (20 μM). The graph on the right shows densitometric analysis of the fold intensity of TGF-βRI, normalized to the loading control. E. Representative immunoblot showing p62 expression in J774 ^WT^ and J774 ^Coro1A^ ^KD^ cells with and without Bafilomycin-A treatment (100 nM). F. Representative immunoblot showing poly-ubiquitination in J774 ^WT^ and J774 ^Coro1A^ ^KD^ cells with and without MG-132 treatment (20 μM). Each dot represents (C and D) one independent experiment of three biological replicates. Actin is used as a loading control (A, D, E, and F). The immunoblots shown here are representative of three biological replicates. Error bars represent mean ± SEM. Statistical analysis performed using the unpaired Student’s t-test, with a p-value <0.05 considered statistically significant. *<0.05, **<0.01, ***< 0.001, ****<0.0001, ns: non-significant.

**Supplementary Figure 7.**
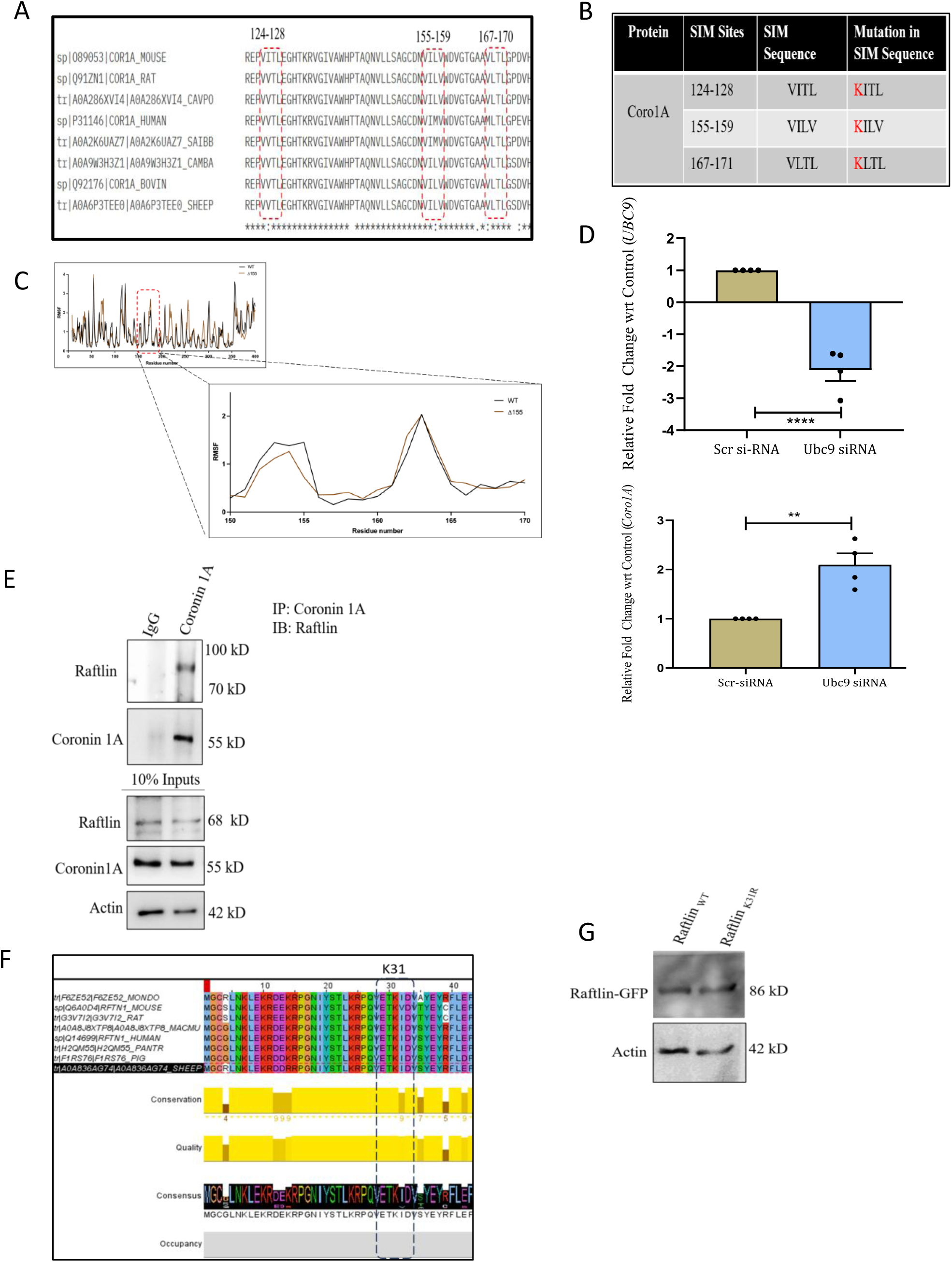
A. Multiple sequence alignment of Coro1A protein sequences from eight mammalian species reveals that selected SIM sites are conserved (highlighted in a red rectangle). B. Table representing site-directed mutations performed in SUMO-interacting motifs (SIM) sites; Valine (V) of the SIM site replaced by Lysine (K). C. RMSF plots of Coro1A^WT^ and Coro1A^V155K^ mutant showing fluctuations of residues observed in molecular dynamics run by the CABS flex server. D. qRT-PCR-based fold change expression of UBC9 and Coro1A from RAW264.7 cells transfected with Scr siRNA and UBC9 siRNA. GAPDH was used for gene normalization. Each dot represents one independent experiment. E. Top: RAW 264.7 cell lysates immunoprecipitated with anti-Coro1A antibody and immunoblotted for Raftlin. Bottom:10% of input loaded from the same cell lysate and immunoblotted for Coro1A, Raftlin, and Actin. Isotype antibodies (IgG) were used as a negative control. F. Multiple sequence alignment of Raftlin protein sequences from eight mammalian species reveals a conserved SUMO-binding site (K31) (highlighted in a black rectangle). G. Representative immunoblot showing stable expression of Raftlin ^K31R^, similar to Raftlin ^WT^.

## References

1. Foersch S, Neurath MF. Colitis-associated neoplasia: Molecular basis and clinical translation. Vol. 71, Cellular and Molecular Life Sciences Birkhauser Verlag AG; 2014. p. 3523–35.

2. Zhou RW, Harpaz N, Itzkowitz SH, Parsons RE. Molecular mechanisms in colitis-associated colorectal cancer. Vol. 12, Oncogenesis Springer Nature; 2023.

3. Díaz-Gay M, dos Santos W, Moody S, Kazachkova M, Abbasi A, Steele CD, et al. Geographic and age variations in mutational processes in colorectal cancer. Nature. Nature Research; 2025;643(8070):230–40. DOI: 10.1038/s41586-025-09025-8

4. Bogaert J, Prenen H. Molecular genetics of colorectal cancer. Vol. 27, Annals of Gastroenterology 2014. p. 9–14.

5. Su Z, El Hage M, Linnebacher M. Mutation patterns in colorectal cancer and their relationship with prognosis. Heliyon. Elsevier Ltd; 2024;10(17). DOI: 10.1016/j.heliyon.2024.e36550

6. Yin Y, Wan J, Yu J, Wu K. Molecular Pathogenesis of Colitis-associated Colorectal Cancer: Immunity, Genetics, and Intestinal Microecology. Vol. 29, Inflammatory Bowel Diseases Oxford University Press; 2023. p. 1648–57.

7. Yamagishi H, Kuroda H, Imai Y, Hiraishi H. Molecular pathogenesis of sporadic colorectal cancers. Vol. 35, Chinese Journal of Cancer Landes Bioscience; 2016.

8. Yashiro M. Ulcerative colitis-associated colorectal cancer. World J Gastroenterol. WJG Press; 2014;20(44):16389–97. DOI: 10.3748/wjg.v20.i44.16389

9. Shawki S, Ashburn J, Signs SA, Huang E. Colon Cancer: Inflammation-Associated Cancer. Vol. 27, Surgical Oncology Clinics of North America W.B. Saunders; 2018. p. 269–87.

10. Kakiuchi N, Yoshida K, Uchino M, Kihara T, Akaki K, Inoue Y, et al. Frequent mutations that converge on the NFKBIZ pathway in ulcerative colitis. Nature. Nature Research; 2020;577(7789):260–5. DOI: 10.1038/s41586-019-1856-1

11. Dan WY, Zhou GZ, Peng LH, Pan F. Update and latest advances in mechanisms and management of colitis-associated colorectal cancer. Vol. 15, World Journal of Gastrointestinal Oncology Baishideng Publishing Group Inc; 2023. p. 1317–31.

12. Deng Y, Jia X, Liu L, He Q, Liu L. The role of intestinal macrophage polarization in colitis-associated colon cancer. Vol. 16, Frontiers in Immunology Frontiers Media SA; 2025.

13. Zhang M, Li X, Zhang Q, Yang J, Liu G. Roles of macrophages on ulcerative colitis and colitis-associated colorectal cancer. Vol. 14, Frontiers in Immunology Frontiers Media S.A.; 2023.

14. Shin AE, Tesfagiorgis Y, Larsen F, Derouet M, Zeng PYF, Good HJ, et al. F4/80+Ly6Chigh Macrophages Lead to Cell Plasticity and Cancer Initiation in Colitis. Gastroenterology. W.B. Saunders; 2023;164(4):593–609.e13. DOI: 10.1053/j.gastro.2023.01.002

15. Hardbower DM, Coburn LA, Asim M, Singh K, Sierra JC, Barry DP, et al. EGFR-mediated macrophage activation promotes colitis-associated tumorigenesis. Oncogene. Nature Publishing Group; 2017;36(27):3807–19. DOI: 10.1038/onc.2017.23

16. Wang Y, Liu H, Zhang Z, Bian D, Shao K, Wang S, et al. G-MDSC-derived exosomes mediate the differentiation of M-MDSC into M2 macrophages promoting colitis-to-cancer transition. J Immunother Cancer. BMJ Publishing Group; 2023;11(6). DOI: 10.1136/jitc-2022-006166

17. Yuan Q, Gu J, Zhang J, Liu S, Wang Q, Tian T, et al. MyD88 in myofibroblasts enhances colitis-associated tumorigenesis via promoting macrophage M2 polarization. Cell Rep. Elsevier B.V.; 2021;34(5). DOI: 10.1016/j.celrep.2021.108724

18. Sheng YH, Davies JM, Wang R, Wong KY, Giri R, Yang Y, et al. MUC1-mediated Macrophage Activation Promotes Colitis-associated Colorectal Cancer via Activating the Interleukin-6/ Signal Transducer and Activator of Transcription 3 Axis. CMGH. Elsevier Inc.; 2022;14(4):789–811. DOI: 10.1016/j.jcmgh.2022.06.010

19. Long Y, Huang F, Zhang J, Zhang J, Cheng R, Zhu L, et al. Identification of SUMOylation-related signature genes associated with immune infiltration in ulcerative colitis through bioinformatics analysis and experimental validation. Gene. Elsevier B.V.; 2025;935. DOI: 10.1016/j.gene.2024.148996

20. Ma XN, Li MY, Qi GQ, Wei LN, Zhang DK. SUMOylation at the crossroads of gut health: insights into physiology and pathology. Vol. 22, Cell Communication and Signaling BioMed Central Ltd; 2024.

21. Gostissa M, Hengstermann A, Fogal V, Sandy P, Schwarz SE, Scheffner M, et al. Activation of p53 by conjugation to the ubiquitin-like protein SUMO-1. Vol. 18, The EMBO Journal 1999.

22. Zhou Z, Wang M, Li J, Xiao M, Chin YE, Cheng J, et al. SUMOylation and SENP3 regulate STAT3 activation in head and neck cancer. Oncogene. Nature Publishing Group; 2016;35(45):5826–38. DOI: 10.1038/onc.2016.124

23. Yang Q, Tang J, Xu C, Zhao H, Zhou Y, Wang Y, et al. Histone deacetylase 4 inhibits NF-κB activation by facilitating IκBα sumoylation. J Mol Cell Biol. Oxford University Press; 2020;12(12):933–45. DOI: 10.1093/jmcb/mjaa043

24. Lin CH, Liu SY, Lee EHY. SUMO modification of Akt regulates global SUMOylation and substrate SUMOylation specificity through Akt phosphorylation of Ubc9 and SUMO1. Oncogene. Nature Publishing Group; 2016;35(5):595–607. DOI: 10.1038/onc.2015.115

25. Liu T, Wang H, Chen Y, Wan Z, Du Z, Shen H, et al. SENP5 promotes homologous recombination-mediated DNA damage repair in colorectal cancer cells through H2AZ deSUMOylation. Journal of Experimental and Clinical Cancer Research. BioMed Central Ltd; 2023;42(1). DOI: 10.1186/s13046-023-02789-9

26. Sheng Z, Luo S, Huang L, Deng YN, Zhang N, Luo Y, et al. SENP1-mediated deSUMOylation of YBX1 promotes colorectal cancer development through the SENP1-YBX1-AKT signaling axis. Oncogene. Springer Nature; 2025;44(19):1361–74. DOI: 10.1038/s41388-025-03302-6

27. Du L, Li YJ, Fakih M, Wiatrek RL, Duldulao M, Chen Z, et al. Role of SUMO activating enzyme in cancer stem cell maintenance and self-renewal. Nat Commun. Nature Publishing Group; 2016;7. DOI: 10.1038/ncomms12326

28. Arnesen H, Müller MHB, Aleksandersen M, Østby GC, Carlsen H, Paulsen JE, et al. Induction of colorectal carcinogenesis in the C57BL/6J and A/J mouse strains with a reduced DSS dose in the AOM/DSS model. Lab Anim Res. BioMed Central Ltd; 2021;37(1). DOI: 10.1186/s42826-021-00096-y

29. Ren J, Sui H, Fang F, Li Q, Li B. The application of ApcMin/+ mouse model in colorectal tumor researches. Vol. 145, Journal of Cancer Research and Clinical Oncology Springer Science and Business Media Deutschland GmbH; 2019. p. 1111–22.

30. Sánchez-Alba L, Ying L, Maletic MD, De Bolòs A, Borràs-Gas H, Liu B, et al. Structural basis for the human SENP5’s SUMO isoform discrimination. Nature Communications. Nature Research; 2025;16(1). DOI: 10.1038/s41467-025-60029-4

31. Galkin VE, Orlova A, Brieher W, Yuan Kueh H, Mitchison TJ, Egelman EH. Author manuscript. Vol. 376, J Mol Biol 2008.

32. Humphries CL, Balcer HI, D’Agostino JL, Winsor B, Drubin DG, Barnes G, et al. Direct regulation of Arp2/3 complex activity and function by the actin binding protein coronin. Journal of Cell Biology. 2002;159(6):993–1004. DOI: 10.1083/jcb.200206113

33. Mueller P, Massner J, Jayachandran R, Combaluzier B, Albrecht I, Gatfield J, et al. Regulation of T cell survival through coronin-1-mediated generation of inositol-1,4,5-trisphosphate and calcium mobilization after T cell receptor triggering. Nat Immunol. 2008;9(4):424–31. DOI: 10.1038/ni1570

34. Stocker TJ, Pircher J, Skenderi A, Ehrlich A, Eberle C, Megens RTA, et al. The Actin Regulator Coronin-1A Modulates Platelet Shape Change and Consolidates Arterial Thrombosis. Thromb Haemost. Georg Thieme Verlag; 2018;118(12):2098–111. DOI: 10.1055/s-0038-1675604

35. Castro-Castro A, Ojeda V, Barreira M, Sauzeau V, Navarro-Lérida I, Muriel O, et al. Coronin 1A promotes a cytoskeletal-based feedback loop that facilitates Rac1 translocation and activation. EMBO Journal. 2011;30(19):3913–27. DOI: 10.1038/emboj.2011.310

36. Pick R, Begandt D, Stocker TJ, Salvermoser M, Thome S, Böttcher RT, et al. Coronin 1A, a novel player in integrin biology, controls neutrophil trafficking in innate immunity. Blood. American Society of Hematology; 2017;130(7):847–58. DOI: 10.1182/blood-2016-11-749622

37. Nicolaou O, Sokratous K, Makowska Z, Morell M, De Groof A, Montigny P, et al. Proteomic analysis in lupus mice identifies Coronin-1A as a potential biomarker for lupus nephritis. Arthritis Res Ther. BioMed Central; 2020;22(1). DOI: 10.1186/s13075-020-02236-6

38. Siegmund K, Klepsch V, Hermann-Kleiter N, Baier G. Proof of principle for a T lymphocyte intrinsic function of Coronin 1A. Journal of Biological Chemistry. American Society for Biochemistry and Molecular Biology Inc.; 2016;291(42):22086–92. DOI: 10.1074/jbc.M116.748012

40. Deng X, Liu B, Jiang Q, Li G, Li J, Xu K. CREBH promotes autophagy to ameliorate NASH by regulating Coro1a. Biochim Biophys Acta Mol Basis Dis. Elsevier B.V.; 2024;1870(1). DOI: 10.1016/j.bbadis.2023.166914

41. Gunasekera DC, Ma J, Vacharathit V, Shah P, Ramakrishnan A, Uprety P, et al. The development of colitis in Il10 −/− mice is dependent on IL-22. Mucosal Immunol. Springer Nature; 2020;13(3):493–506. DOI: 10.1038/s41385-019-0252-3

42. Suhail A, Rizvi ZA, Mujagond P, Ali SA, Gaur P, Singh M, et al. DeSUMOylase SENP7-Mediated Epithelial Signaling Triggers Intestinal Inflammation via Expansion of Gamma-Delta T Cells. Cell Rep. Elsevier B.V.; 2019;29(11):3522–3538.e7. DOI: 10.1016/j.celrep.2019.11.028

43. Kaminski S, Hermann-Kleiter N, Meisel M, Thuille N, Cronin S, Hara H, et al. Coronin 1A is an essential regulator of the TGFβ receptor/SMAD3 signaling pathway in Th17 CD4 + T cells. J Autoimmun. 2011;37(3):198–208. DOI: 10.1016/j.jaut.2011.05.018

44. Burandt E, Lübbersmeyer F, Gorbokon N, Büscheck F, Luebke AM, Menz A, et al. E-Cadherin expression in human tumors: a tissue microarray study on 10,851 tumors. Biomark Res. BioMed Central Ltd; 2021;9(1). DOI: 10.1186/s40364-021-00299-4

45. Rodriguez FJ, Lewis-Tuffin LJ, Anastasiadis PZ. E-cadherin’s dark side: Possible role in tumor progression. Vol. 1826, Biochimica et Biophysica Acta - Reviews on Cancer 2012. p. 23–31.

46. Popp C, Nichita L, Voiosu T, Bastian A, Cioplea M, Micu G, et al. Expression Profile of p53 and p21 in Large Bowel Mucosa as Biomarkers of Inflammatory-Related Carcinogenesis in Ulcerative Colitis. Dis Markers. Hindawi Limited; 2016;2016. DOI: 10.1155/2016/3625279

47. Lu X, Yu Y, Tan S. p53 expression in patients with ulcerative colitis - associated with dysplasia and carcinoma: A systematic meta-analysis. BMC Gastroenterol. BioMed Central Ltd.; 2017;17(1). DOI: 10.1186/s12876-017-0665-y

48. Khursheed S, Ali T, Mushtaq M, Humayun S, Khan A, Akbar A, et al. Immunohistochemical Analysis of the p53 Protein in Colorectal Cancer: A Clinicopathological Study. Cureus. Springer Science and Business Media LLC; 2024; DOI: 10.7759/cureus.76172

49. Suarez-Lopez L, Sriram G, Kong YW, Morandell S, Merrick KA, Hernandez Y, et al. MK2 contributes to tumor progression by promoting M2 macrophage polarization and tumor angiogenesis. Proc Natl Acad Sci U S A. National Academy of Sciences; 2018;115(18):E4236–44. DOI: 10.1073/pnas.1722020115

50. Zhang F, Wang H, Wang X, Jiang G, Liu H, Zhang G, et al. TGF-β induces M2-like macrophage polarization via SNAIL-mediated suppression of a pro-inflammatory phenotype [Internet]. Vol. 7 2016 [cited. Available from: www.impactjournals.com/oncotarget

51. Deng Z, Fan T, Xiao C, Tian H, Zheng Y, Li C, et al. TGF-β signaling in health, disease, and therapeutics. Vol. 9, Signal Transduction and Targeted Therapy Springer Nature; 2024.

52. Massagué J. TGFβ signalling in context. Vol. 13, Nature Reviews Molecular Cell Biology 2012. p. 616–30.

53. Jayachandran R, Sundaramurthy V, Combaluzier B, Mueller P, Korf H, Huygen K, et al. Survival of Mycobacteria in Macrophages Is Mediated by Coronin 1-Dependent Activation of Calcineurin. Cell. Elsevier B.V.; 2007;130(1):37–50. DOI: 10.1016/j.cell.2007.04.043

54. Suo D, Park J, Harrington AW, Zweifel LS, Mihalas S, Deppmann CD. Coronin-1 is a neurotrophin endosomal effector that is required for developmental competition for survival. Nat Neurosci. 2014;17(1):36–45. DOI: 10.1038/nn.3593

55. Huang W, Ghisletti S, Saijo K, Gandhi M, Aouadi M, Tesz GJ, et al. Coronin 2A mediates actin-dependent de-repression of inflammatory response genes. Nature. 2011;470(7334):414–8. DOI: 10.1038/nature09703

56. Kerscher O. SUMO junction - What’s your function? New insights through SUMO-interacting motifs. Vol. 8, EMBO Reports 2007. p. 550–5.

57. Song J, Durrin LK, Wilkinson TA, Krontiris TG, Chen Y. Identification of a SUMO-binding motif that recognizes SUMO-modified proteins [Internet]. Vol. 101 2004 [cited. Available from: www.pnas.orgcgidoi10.1073pnas.0403498101

58. Gou Y, Liu D, Chen M, Wei Y, Huang X, Han C, et al. GPS-SUMO 2.0: an updated online service for the prediction of SUMOylation sites and SUMO-interacting motifs. Nucleic Acids Res. Oxford University Press; 2024;52(W1):W238–47. DOI: 10.1093/nar/gkae346

59. Beauclair G, Bridier-Nahmias A, Zagury JF, Säb A, Zamborlini A. JASSA: A comprehensive tool for prediction of SUMOylation sites and SIMs. Bioinformatics. Oxford University Press; 2015;31(21):3483–91. DOI: 10.1093/bioinformatics/btv403

60. Rytinki MM, Palvimo JJ. SUMOylation attenuates the function of PGC-1α. Journal of Biological Chemistry. 2009;284(38):26184–93. DOI: 10.1074/jbc.M109.038943

61. Niu X, Xu C, Cheuk YC, Xu X, Liang L, Zhang P, et al. Characterizing hub biomarkers for post-transplant renal fibrosis and unveiling their immunological functions through RNA sequencing and advanced machine learning techniques. J Transl Med. BioMed Central Ltd; 2024;22(1). DOI: 10.1186/s12967-024-04971-9

62. Elihamu D, Li Y, Wang Y, Cui H, Xing Y, Peng H, et al. CORO1A: a pan-cancer prognosis, diagnostic and immune biomarker based on breast cancer validation. Front Oncol. Frontiers Media SA; 2025;15. DOI: 10.3389/fonc.2025.1670526

63. Kros JM, Zeneyedpour L, Pedrosa RMSM, Belcaid Z, Dik WA, Luider TM, et al. T cell induced expression of Coronin-1A facilitates blood-brain barrier transmigration of breast cancer cells. Sci Rep. Nature Research; 2024;14(1). DOI: 10.1038/s41598-024-83301-x

64. Li Z, Chen L, Wei Z, Liu H, Zhang L, Huang F, et al. A novel classification method for LUAD that guides personalized immunotherapy on the basis of the cross-talk of coagulation- and macrophage-related genes. Front Immunol. Frontiers Media SA; 2025;16. DOI: 10.3389/fimmu.2025.1518102

65. Sun Y, Ma Q, Chen Y, Liao D, Kong F. Identification and analysis of prognostic immune cell homeostasis characteristics in lung adenocarcinoma. Clinical Respiratory Journal. John Wiley and Sons Inc; 2024;18(5). DOI: 10.1111/crj.13755

66. Zhou Y, Shi X, Chen H, Mao B, Song X, Gao L, et al. Tumor Immune Microenvironment Characterization of Primary Lung Adenocarcinoma and Lymph Node Metastases. Biomed Res Int. Hindawi Limited; 2021;2021. DOI: 10.1155/2021/5557649

67. Bao M, Huang Y, Lang Z, Zhao H, Saito Y, Nagano T, et al. Proteomic analysis of plasma exosomes in patients with non-small cell lung cancer. Transl Lung Cancer Res. AME Publishing Company; 2022;11(7):1434–52. DOI: 10.21037/tlcr-22-467

68. Kumar P, Nandi S, Tan TZ, Ghee Ler S, Chia KS, Lim W-Y, et al. Highly sensitive and specific novel biomarkers for the diagnosis of transitional bladder carcinoma [Internet]. Available from: www.impactjournals.com/oncotarget/

69. Jayachandran R, Gumienny A, Bolinger B, Ruehl S, Lang MJ, Fucile G, et al. Disruption of Coronin 1 Signaling in T Cells Promotes Allograft Tolerance while Maintaining Anti-Pathogen Immunity. Immunity. Cell Press; 2019;50(1):152–165.e8. DOI: 10.1016/j.immuni.2018.12.011

70. Ndinyanka Fabrice T, Buczak K, Schmidt A, Pieters J. T cell population size control by coronin 1 uncovered: from a spot identified by two-dimensional gel electrophoresis to quantitative proteomics. Vol. 22, Expert Review of Proteomics Taylor and Francis Ltd.; 2025. p. 35–44.

71. Jayachandran R, Pieters J. Regulation of immune cell homeostasis and function by coronin 1. Vol. 28, International Immunopharmacology Elsevier B.V.; 2015. p. 825–8.

72. Shiow LR, Paris K, Akana MC, Cyster JG, Sorensen RU, Puck JM. Severe combined immunodeficiency (SCID) and attention deficit hyperactivity disorder (ADHD) associated with a coronin-1A mutation and a chromosome 16p11.2 deletion. Clinical Immunology. 2009;131(1):24–30. DOI: 10.1016/j.clim.2008.11.002

73. Younis DA, Marosvari M, Liu W, Pulikkot S, Cao Z, Zhou B, et al. CFTR dictates monocyte adhesion by facilitating integrin clustering but not activation. Proc Natl Acad Sci U S A. National Academy of Sciences; 2025;122(3). DOI: 10.1073/pnas.2412717122

74. Deng Z, Fan T, Xiao C, Tian H, Zheng Y, Li C, et al. TGF-β signaling in health, disease, and therapeutics. Vol. 9, Signal Transduction and Targeted Therapy Springer Nature; 2024.

75. Fridlender ZG, Sun J, Kim S, Kapoor V, Cheng G, Ling L, et al. Polarization of Tumor-Associated Neutrophil Phenotype by TGF-β: “N1” versus “N2” TAN. Cancer Cell. Cell Press; 2009;16(3):183–94. DOI: 10.1016/j.ccr.2009.06.017

76. Gong D, Shi W, Yi S ju, Chen H, Groffen J, Heisterkamp N. TGFβ signaling plays a critical role in promoting alternative macrophage activation. BMC Immunol. 2012;13. DOI: 10.1186/1471-2172-13-31

77. Lin CS, Chen MF, Wang YS, Chuang TF, Chiang YL, Chu RM. IL-6 restores dendritic cell maturation inhibited by tumor-derived TGF-β through interfering Smad 2/3 nuclear translocation. Cytokine. 2013;62(3):352–9. DOI: 10.1016/j.cyto.2013.03.005

78. Mckarns SC, Schwartz RH, Kaminski NE. Smad3 Is Essential for TGF-1 to Suppress IL-2 Production and TCR-Induced Proliferation, but Not IL-2-Induced Proliferation1 [Internet]. 2004 [cited. Available from: https://academic.oup.com/jimmunol/article/172/7/4275/8060313

79. Strobl H, Knapp W. TGF-1 regulation of dendritic cells TGF-/ dendritic cell / Langerhans cell / differentiation / antigen presentation. 1999.

80. Calon A, Espinet E, Palomo-Ponce S, Tauriello DVF, Iglesias M, Céspedes MV, et al. Dependency of Colorectal Cancer on a TGF-β-Driven Program in Stromal Cells for Metastasis Initiation. Cancer Cell. Cell Press; 2012;22(5):571–84. DOI: 10.1016/j.ccr.2012.08.013

81. Means AL, Freeman TJ, Zhu J, Woodbury LG, Marincola-Smith P, Wu C, et al. Epithelial Smad4 Deletion Up-Regulates Inflammation and Promotes Inflammation-Associated Cancer. CMGH. Elsevier Inc; 2018;6(3):257–76. DOI: 10.1016/j.jcmgh.2018.05.006

82. Gong D, Shi W, Yi S ju, Chen H, Groffen J, Heisterkamp N. TGFβ signaling plays a critical role in promoting alternative macrophage activation. BMC Immunol. 2012;13. DOI: 10.1186/1471-2172-13-31

83. Chan MKK, Chung JYF, Tang PCT, Chan ASW, Ho JYY, Lin TPT, et al. TGF-β signaling networks in the tumor microenvironment. Vol. 550, Cancer Letters Elsevier Ireland Ltd; 2022.

84. Li J, Liu Y, Wang B, Xu Y, Ma A, Zhang F, et al. Myeloid TGF-β signaling contributes to colitis-associated tumorigenesis in mice. Carcinogenesis. 2013;34(9):2099–108. DOI: 10.1093/carcin/bgt172

85. Wang W, Li X, Zheng D, Zhang D, Peng X, Zhang X, et al. Dynamic changes and functions of macrophages and M1/M2 subpopulations during ulcerative colitis-associated carcinogenesis in an AOM/DSS mouse model. Mol Med Rep. Spandidos Publications; 2015;11(4):2397–406. DOI: 10.3892/mmr.2014.3018

86. Zadka Ł, Chabowski M, Grybowski D, Piotrowska A, Dzięgiel P. Interplay of stromal tumor-infiltrating lymphocytes, normal colonic mucosa, cancer-associated fibroblasts, clinicopathological data and the immunoregulatory molecules of patients diagnosed with colorectal cancer. Cancer Immunology, Immunotherapy. Springer Science and Business Media Deutschland GmbH; 2021;70(9):2681–700. DOI: 10.1007/s00262-021-02863-1

87. Kazuko Saeki, Akihiko Yoshimura. The B-cell-specific major raft protein, Raftlin, is necessary for the integrity of lipid raft and BCR signal transduction. EMBO. 2003;12:3015–26.

88. Saeki K, Fukuyama S, Ayada T, Nakaya M, Aki D, Takaesu G, et al. A Major Lipid Raft Protein Raftlin Modulates T Cell Receptor Signaling and Enhances Th17-Mediated Autoimmune Responses. The Journal of Immunology. Oxford University Press (OUP); 2009;182(10):5929–37. DOI: 10.4049/jimmunol.0802672

89. Watanabe A, Tatematsu M, Saeki K, Shibata S, Shime H, Yoshimura A, et al. Raftlin is involved in the nucleocapture complex to induce poly(I:C)-mediated TLR3 activation. Journal of Biological Chemistry. American Society for Biochemistry and Molecular Biology Inc.; 2011;286(12):10702–11. DOI: 10.1074/jbc.M110.185793

90. Bayliss AL, Sundararaman A, Granet C, Mellor H. Raftlin is recruited by neuropilin-1 to the activated VEGFR2 complex to control proangiogenic signaling. Angiogenesis. Springer; 2020;23(3):371–83. DOI: 10.1007/s10456-020-09715-z

91. Tatematsu M, Yoshida R, Morioka Y, Ishii N, Funami K, Watanabe A, et al. Raftlin Controls Lipopolysaccharide-Induced TLR4 Internalization and TICAM-1 Signaling in a Cell Type–Specific Manner. The Journal of Immunology. Oxford University Press (OUP); 2016;196(9):3865–76. DOI: 10.4049/jimmunol.1501734

92. Lacy MM, Ma R, Ravindra NG, Berro J. Molecular mechanisms of force production in clathrin-mediated endocytosis. Vol. 592, FEBS Letters Wiley Blackwell; 2018. p. 3586–605.

93. Mustfa SA, Singh M, Suhail A, Mohapatra G, Verma S, Chakravorty D, et al. SUMOylation pathway alteration coupled with downregulation of SUMO E2 enzyme at mucosal epithelium modulates inflammation in inflammatory bowel disease. Open Biol. Royal Society Publishing; 2017;7(6). DOI: 10.1098/rsob.170024

94. Toda G, Yamauchi T, Kadowaki T, Ueki K. Preparation and culture of bone marrow-derived macrophages from mice for functional analysis. STAR Protoc. Cell Press; 2021;2(1). DOI: 10.1016/j.xpro.2020.100246

95. Weigmann B, Tubbe I, Seidel D, Nicolaev A, Becker C, Neurath MF. Isolation and subsequent analysis of murine lamina propria mononuclear cells from colonic tissue. Nat Protoc. 2007;2(10):2307–11. DOI: 10.1038/nprot.2007.315

